# The role of cortico-thalamic feedback for visual information processing

**DOI:** 10.1101/2025.01.27.635001

**Authors:** Wenqing Wei, Stefan Rotter

## Abstract

In the early visual system, neurons in the dorsal lateral geniculate nucleus (dLGN) not only relay visual inputs from the retina to the primary visual cortex (V1), but also receive cortico-thalamic (CT) feedback inputs, which anatomically outnumber feedforward inputs. Most experimental and theoretical work so far has focused on the modulatory influence of CT feedback on relay cells in the dLGN. However, the role of feedback for the activity of V1 neurons, which are mainly driven by visual input meditated by dLGN neurons, is still unclear. To address this question, we devised a spiking neural network model of the thalamo-cortical and cortico-thalamic loop, where the CT feedback can be turned on and off to study its effect. We identified three main phenomena of the effect of CT feedback on V1 neurons: (i) orientation selectivity of cortical responses is increased for all contrasts, (ii) preferred orientations of input and output are better aligned with each other, (iii) neuronal population activity represents the stimulus orientation with higher confidence. Our analysis showed that CT feedback generally improves the signal-to-noise ratio (SNR) of the responses of dLGN neurons and therefore also increases the SNR of thalamic inputs to V1 neurons. Moreover, we found that the modulatory effects on visual processing depend on the geometry of the CT feedback projections. We conclude that CT feedback increases the selectivity of V1 neurons particularly at low contrasts by modulating the activity of dLGN neurons.

**Author summary:** Extensive experimental and theoretical studies have been conducted to understand the functional role of cortico-thalamic (CT) feedback on relay cells in the dorsal lateral geniculate nucleus (dLGN) during visual processing. However, it remains unclear how CT feedback affects the neuronal responses of V1 neurons. To address this question, we developed a computational model of the thalamo-cortico-thalamic loop circuit based on biophysical parameters. Our results indicate that CT feedback modulates the orientation preference of V1 neurons in a contrast-dependent manner: The increase of orientation selectivity (OS) is stronger and the input preferred orientation (PO) is better preserved at lower contrasts. Our analysis suggests a mechanistic explanation: The CT feedback attenuates the untuned part of the thalamic input, while it amplifies the tuned part. This effectively increases the signal-to-noise ratio (SNR) in particular for weak contrasts. Our network model for the first time allows us to study the effect of feedback on cortical neurons and understand the impact of the number and the geometry of feedback connections.

## Introduction

In the mammalian visual system, light-induced signals from retinal ganglion cells (RGC) are transmitted to the primary visual cortex (V1) via thalamo-cortical (TC) relay cells in the dorsal lateral geniculate nucleus (dLGN). However, visual information is not only transmitted via feedforward pathways. Pyramidal cells in layer 6 of V1 establish synapses to relay cells and interneurons in dLGN as well as GABAergic neurons in thalamic reticular nucleus (TRN), which in turn innervates dLGN relay cells [1]. It has been observed in different species that these feedback connections often outnumber feedforward afferents anatomically [2, 3]. What then is the role of cortico-thalamic feedback connections in view of their high biological cost? In contrast to feedforward projections that are thought to drive the visual selective responses [4–6], the cortico-thalamic (CT) feedback is generally considered to be modulatory [7]. Although the effect of feedback on sensory processing was studied in many experiments across several animal species [8–11], a general consensus on the functional role of CT feedback has not been reached.

Since TC cells in dLGN receive both direct excitation from cortex and indirect inhibition via TRN cells and thalamic interneurons, the overall effect of CT feedback is a complex mix. Most experimental and theoretical studies in the past focused on the impact of CT feedback on the activity and receptive fields (RFs) of TC cells. For example, CT feedback has been reported to have an effect on switching the firing pattern of TC cells between burst and tonic modes [12–14], enhancing the center-surround antagonism of the RFs [15–17] and modulating the visual transmission differently under different visual stimuli [14]. However, due to experimental difficulties, less is known about the exact role of CT feedback on V1 neurons with respect to their functional selectivity.

Using orientation selectivity as an example, we seek to characterize the effects of CT feedback on the responses of cortical neurons and identify a possible mechanism via spiking network simulations. Here we developed a computational network model of the thalamo-cortico-thalamic circuit based on our previous work [18], where the orientation selectivity of V1 neurons emerges in the feedforward pathway. In our extended model, we are able to turn the CT feedback on and off and record the neuronal responses at all stages. By comparing the neuronal responses of simulations of the feedforward network with CT feedback turned off, and the network with feedback, we observed a series of significant effects of CT feedback on the orientation preference of V1 neurons. First, the orientation selectivity (OS) of V1 neurons is enhanced when the CT feedback is present. This effect can be established at all stimulus contrasts, but it is more significant for lower contrasts. Importantly, the correspondence between the preferred orientations (PO) of V1 neurons and their respective thalamic inputs is better preserved in the network with feedback. Furthermore, decoding the stimulus orientation on the population level shows that CT feedback improves the confidence of orientation coding by the network significantly. Our simulations show that the CT effects on the orientation preference depend not only on the geometry of the direct and indirect pathways but also the number of feedback connections.

Our analysis suggested a mechanism for the feedback effects on orientation preference. Compared to the feedforward network, the signal-to-noise ratio (SNR) of the TC cell responses is larger in the CT feedback network. This in turn improves the SNR of the compound thalamic inputs to V1 neurons, which is a superposition of the responses of all presynaptic TC cells that converge to one V1 neuron. Consistent with the output of V1 neurons, the increase of the SNR of their thalamic inputs is generally stronger at lower contrasts. In addition, neuronal responses in RF centers were generally enhanced by the CT feedback.

Overall, our findings imply that the CT feedback may play a key role in making the responses of cortical neurons more selective, especially at low contrasts. This is achieved without changing properties of the input, but by modulating the activity of TC neurons. Using a biologically plausible computational model based on networks of spiking neurons, we were able to provide new insights into the neuronal mechanisms that complement our incomplete knowledge from experiments.

## Methods

### Neuron models

#### RGC cells

Retinal ganglion cells (RGCs) have mostly circular antogonistic center-surround receptive fields. Therefore, in our model the receptive fields of RGCs are represented as a classic difference-of-Gaussian (DoG)

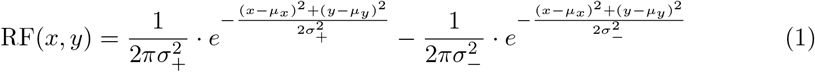

where (*µ*_*x*_, *µ*_*y*_) denotes the position of the receptive field center. For *σ*_+_ < *σ*_−_ we obtain an ON-center cell, for *σ*_+_ *> σ*_−_ we have an OFF-center cell. The time-dependent firing rate *ν*(*t*) of this RGC in response to a visual stimulus is then the spatial convolution of its receptive field and the light intensity of the stimulus *S*(*x, y, t*)

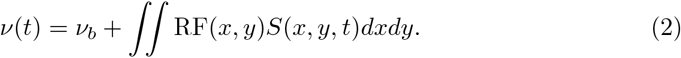

In order to study the orientation selectivity, oriented sinusoidal moving gratings are presented in the entire visual field. The direction of movement of the grating is always perpendicular to its orientation. The light intensity at position (*x, y*) and at time *t* of an oriented grating with orientation *θ*, spatial frequency λ and temporal frequency *f* is specifically given by

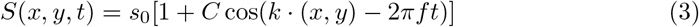

where *s*_0_ is the mean luminance of the stimulus, *k* = 2*π*λ(sin(*θ*), − cos(*θ*)) is the wave vector and *C* ∈ [0, 1] is the contrast of the grating. The time-dependent firing rate of a RGC with RF center at position (*µ*_*x*_, *µ*_*y*_) is then given by

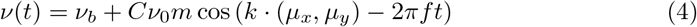

where 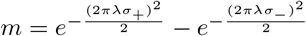 and *ν*_0_ is the rate of the neuron in response to the mean stimulus luminance *s*_0_ and *ν*_*b*_ = *m*_max_*ν*_0_.

The activity of the RGC is conceived as a Poisson process with intensity *ν*(*t*) to reflect the spiking nature of these cells. Individual spikes at time *t*^*′*^ are represented by Dirac delta functions *δ*(*t* − *t*^*′*^). Then the spike train of a neuron is the sum of all its spikes Σ_*k*_ *δ*(*t* − *t*^*k*^).

#### Other neurons

All other neurons in our network except for RGCs are conceived as leaky integrate-and-fire (LIF) neurons, a computational model widely used in the field. The time evolution of the membrane potential of a LIF neuron is determined by the ordinary differential equation

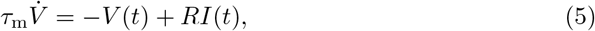

where *τ*_m_ is the membrane time constant and *R* is the leak resistance of the cell. *I*(*t*) is the current reflecting the total synaptic input to the neuron. Assuming that individual post-synaptic currents are scaled delta functions, the synaptic input *I*(*t*) to neuron *i* is given by

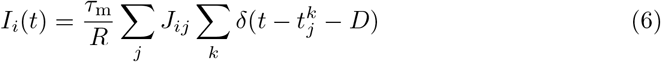

where *J*_*ij*_ represents the amplitude of the postsynaptic potential from presynaptic neuron *j* to its target neuron *i*, and the inner sum reflects the spike train of neuron *j* providing input to neuron *i* with transmission delay *D*. When the input *I*(*t*) is strong enough to depolarize the membrane potential so that it reaches the threshold *V*_th_, an action potential is elicited. Then the membrane potential will be reset to the potential *V*_*r*_. It remains there for a short refractory period *t*_ref_ in which no spikes will be generated. The parameters used for numerical simulation are listed in Table 1.

**Table 1.**
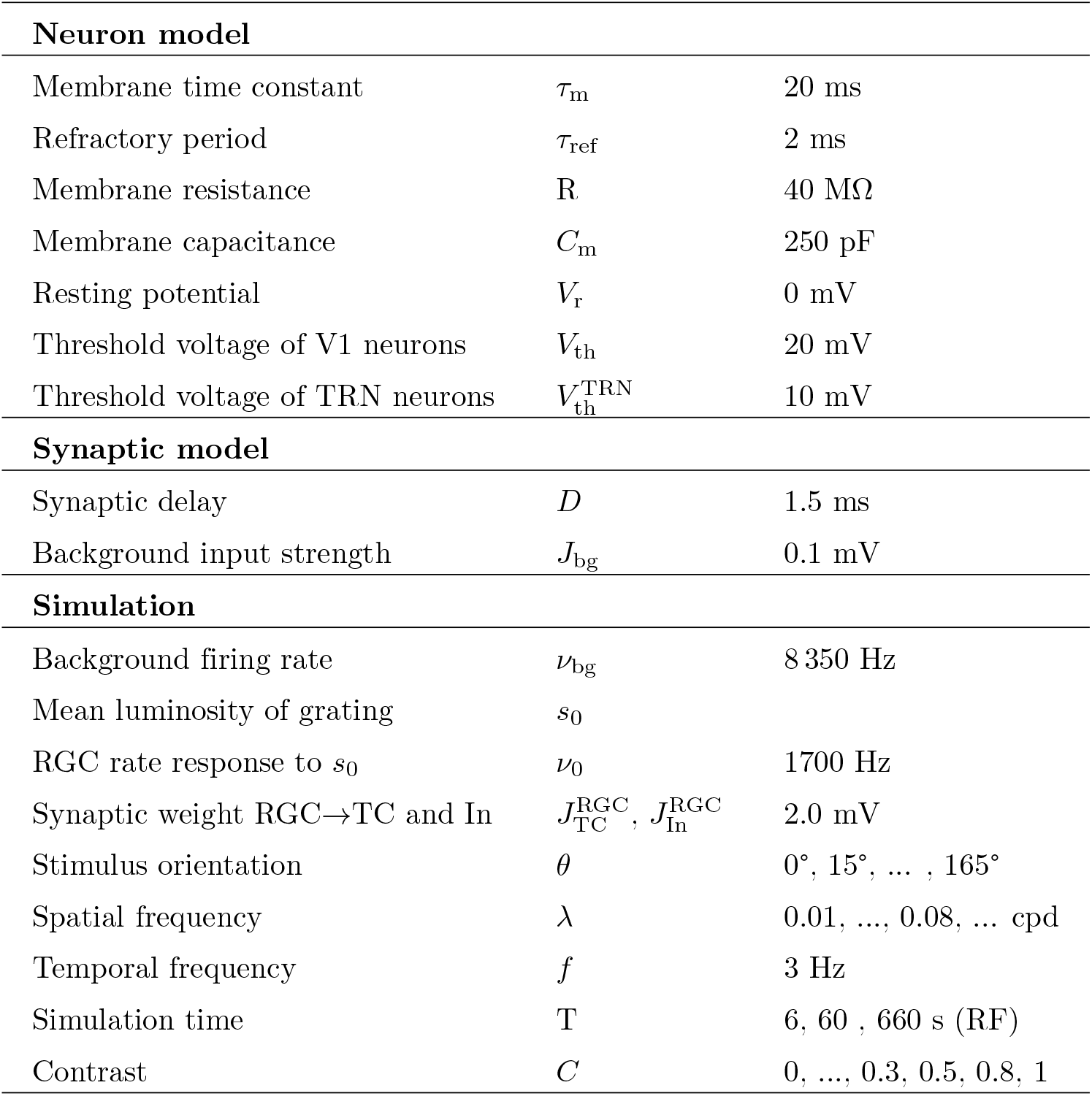
Parameters used for numerical simulations.

| **Neuron model** |  |  |
| --- | --- | --- |
| Membrane time constant | $\tau_m$ | 20 ms |
| Refractory period | $\tau_{\text{ref}}$ | 2 ms |
| Membrane resistance | $R$ | 40 MΩ |
| Membrane capacitance | $C_m$ | 250 pF |
| Resting potential | $V_r$ | 0 mV |
| Threshold voltage of V1 neurons | $V_{\text{th}}$ | 20 mV |
| Threshold voltage of TRN neurons | $V_{\text{th}}^{\text{TRN}}$ | 10 mV |
| **Synaptic model** |  |  |
| Synaptic delay | $D$ | 1.5 ms |
| Background input strength | $J_{\text{bg}}$ | 0.1 mV |
| **Simulation** |  |  |
| Background firing rate | $\nu_{\text{bg}}$ | 8 350 Hz |
| Mean luminosity of grating | $s_0$ | |
| RGC rate response to $s_0$ | $\nu_0$ | 1700 Hz |
| Synaptic weight RGC→TC and In | $J_{\text{TC}}^{\text{RGC}}, J_{\text{In}}^{\text{RGC}}$ | 2.0 mV |
| Stimulus orientation | $\theta$ | 0°, 15°, ... , 165° |
| Spatial frequency | $\lambda$ | 0.01, ..., 0.08, ... cpd |
| Temporal frequency | $f$ | 3 Hz |
| Simulation time | $T$ | 6, 60 , 660 s (RF) |
| Contrast | $C$ | 0, ..., 0.3, 0.5, 0.8, 1 |

### Network configuration

For our study, we have devised a current-based spiking neural network consisting of four parts: retinal ganglion cells (RGC), thalamus, primary visual cortex (V1) and background neurons (see Fig 1). In this network, the cortico-thalamic (CT) feedback can be easily turned on or off to study its functional effects: When the number of CT neurons in the cortex is set to 0, the CT feedback is turned off and the thalamo-cortico-thalamic feedback network becomes the thalamo-cortical feedforward network without CT feedback.

**Fig 1.**
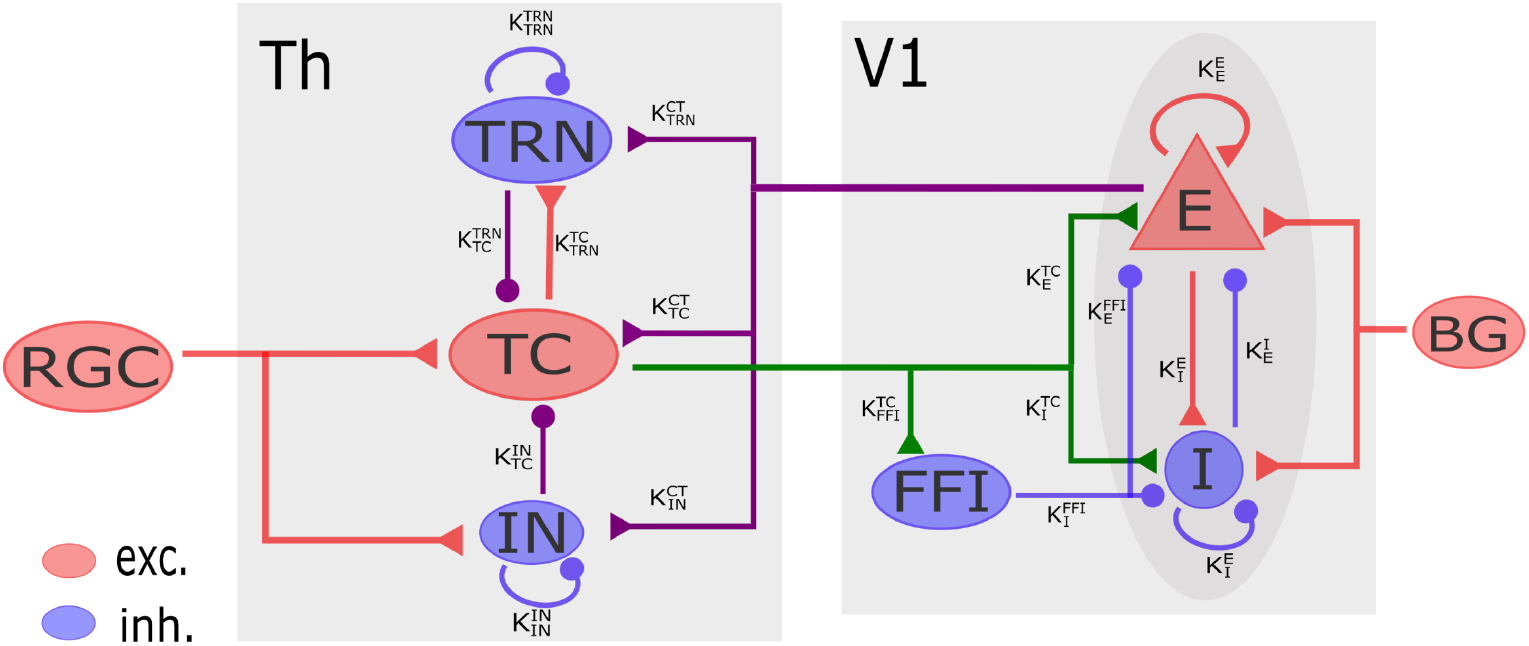
Schematic of the network model. The model is composed of 4 parts. The RGC receives visual input and forwards it to TC cells and interneurons in the thalamus. The thalamus (Th) is comprised of the thalamic reticular nucleus (TRN), thalamo-cortical relay cells (TC) and interneurons (IN). The cortex is represented by a recurrent network with excitatory and inhibitory V1 cells as well as feedforward inhibitory cells (FFI). The background input (BG) accounts for inputs from other parts of the brain. Lines with triangles indicate excitatory projections, dots represent inhibitory connections. The thalamo-cortical feedforward pathways are marked green, cortico-thalamic feedback pathways are purple. In our simulations, the CT feedback can be turned off to study its effect on the rest of the network.

In our model, the RGCs respond to a visual stimulus and further transfer the visual signals to thalamo-cortical relay cells (TC) and interneurons in the dorsal lateral geniculate nucleus (dLGN). The population sizes and connectivity are summarized in Table 2.

**Table 2.** Population sizes and synaptic connections in our model.

|  |  | Pre |  |  |  |  |  |
| --- | --- | --- | --- | --- | --- | --- | --- |
|  |  | TC | TRN | In | FFI | V1<sub>E</sub> | V1<sub>I</sub> |
| population size |  | 3 071 | 2 256 | 300 | 1 500 | 10 000 (incl. CT) | 2 500 |
| Post | TC | 0 | $K_{TC}^{TRN} = 80$ | $K_{TC}^{In} = 5$ | 0 | $K_{TC}^{CT} = 6$ | 0 |
| | | 0 | $J_{TC}^{TRN} = -0.2 \text{ mV}$ | $J_{TC}^{In} = 0.1$ | 0 | $J_{TC}^{CT} = 0.545$ | 0 |
| | TRN | $K_{TRN}^{TC} = 65$ | $K_{TRN}^{TRN} = 150$ | 0 | 0 | $K_{TRN}^{CT} = 65$ | 0 |
| | | $J_{TRN}^{TC} = 0.1 \text{ mV}$ | $J_{TRN}^{TRN} = -0.2 \text{ mV}$ | 0 | 0 | $J_{TRN}^{CT} = 0.6 \text{ mV}$ | 0 |
| | In | 0 | 0 | $K_{In}^{In} = 5$ | 0 | $K_{In}^{CT} = 5$ | 0 |
| | | 0 | 0 | $J_{In}^{In} = -0.2 \text{ mV}$ | 0 | $J_{In}^{CT} = 0.2 \text{ mV}$ | 0 |
| | FFI | $K_{FFI}^{TC} = 35$ | 0 | 0 | 0 | 0 | 0 |
| | | $J_{FFI}^{TC} = 2.0 \text{ mV}$ | 0 | 0 | 0 | 0 | 0 |
| | V1<sub>E</sub> | $K_E^{TC} = 100$ | 0 | 0 | $K_E^{FFI} = 320$ | $K_E^E = 1\,000$ | $K_E^I = 250$ |
| | | $J_E^{TC} = 2.0 \text{ mV}$ | 0 | 0 | $J_E^{FFI} = -2.0 \text{ mV}$ | $J_E^E = 0.2 \text{ mV}$ | $J_E^I = -1.6 \text{ mV}$ |
| | V1<sub>I</sub> | $K_I^{TC} = 100$ | 0 | 0 | $K_I^{FFI} = 320$ | $K_I^E = 1\,000$ | $K_I^I = 250$ |
| | | $J_I^{TC} = 2.0 \text{ mV}$ | 0 | 0 | $J_I^{FFI} = -2.0 \text{ mV}$ | $J_I^E = 0.2 \text{ mV}$ | $J_I^I = -1.6 \text{ mV}$ |

#### Thalamus

There are three types of cells in the thalamus: excitatory thalamo-cortical relay cells (TC) and local inhibitory GABAergic cells including dLGN interneurons and the cells of thalamic reticular nucleus (TRN). The TRN is a shell-shaped structure surrounding the thalamus and located between the thalamus and the cortex. In this structure, each TC cell receives inputs from 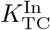 interneurons and projects to 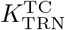 TRN neurons with synaptic weight 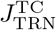. Each TRN neuron in turn makes inhibitory feedback connections onto TC cells with an outdegree of 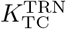with weight 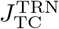. As observed in cats and rats [19–21], TRN neurons also communicate with each other in our model: a TRN neuron receives inputs from 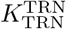TRN neurons with a synaptic efficacy of 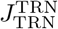.

#### Cortical recurrent network

The layout of the V1 recurrent network is identical to the one introduced by Nicolas Brunel [22]. The recurrent network consists of *N* = 12 500 neurons, of which *a* = 80% are excitatory and 1 − a = 20% are inhibitory. The recurrent connectivity *ϵ* = 10% is the same for all types of neurons in the network [23]. As a result, each neuron receives exactly 1 000 excitatory and 250 inhibitory inputs from within the same network, drawn randomly and independently. Self-connections are excluded. The amplitudes of excitatory recurrent synapses are *J*_E_ = 0.2 mV. Inhibitory couplings are set to be *g* = 8 times stronger than excitatory ones. As a consequence, the amplitudes of inhibitory synapses are *J*_I_ = −*gJ*_E_ = −1.6 mV. This results in an inhibition-dominated recurrent network.

#### Feedforward pathway

Besides the recurrent inputs, V1 neurons receive feedforward inputs from three sources: TC cells, feedforward inhibition and a constant excitatory background. The background inputs represent projections from other parts of the brain except visual thalamus. They are identical for all recurrent neurons to keep the network activity going even in absence of visual stimulation. They are rendered as a stationary Poisson process with constant rate *ν*_bg_ and the synaptic weights are *J*_bg_ = 0.1 mV. The second input source is the visual thalamus. A recurrent V1 neuron receives inputs from 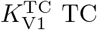 neurons in the thalamus. Experiments *in vitro* reported that thalamo-cortical synapses are several times stronger than intracortical synapses [24, 25]. Here, we assume that the efficacy of direct thalamo-cortical projections is 10 times larger than excitatory recurrent connections, 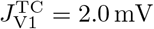. The third source of inputs are feedforward inhibitory projections (FFI) from other cortical neurons. FFI neurons represent a specific type of inhibitory interneurons, which selectively target recurrent V1 neurons. They receive input from 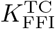 neurons in the thalamus, each with a synaptic efficacy of 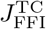. Finally, 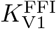 FFI neurons project onto each V1 neuron, each with synaptic weight 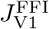. In the cortical circuit just described, the synaptic connections between FFI neurons and recurrent V1 neurons are established randomly and independently.

#### Feedback pathway

In the feedforward pathway, most of the axons from TC cells innervate layer 4 and 6 of the V1 network, while the main source of cortical feedback projections to thalamic neurons originate from layer 6. A complete feedback model will require a micro-circuit in V1 including many parameters which are not fully understood yet. For simplicity, the cortico-thalamic feedback (CT) neurons in our model are chosen randomly from the same group of V1 pyramidal cells that also receive TC inputs.

The CT feedback has two pathways: a direct excitatory pathway and an indirect inhibitory pathway. In the direct feedback pathway, one CT cell directly makes excitatory connections to 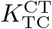 TC cells. In turn, CT neurons inhibit TC cells indirectly through TRN cells and interneurons. In our model, each CT neuron projects to 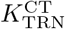 TRN neurons with synaptic weight 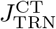 as well as 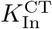 interneurons with weight 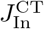. Then these inhibitory neurons further innervate TC cells with the connectivity described in Thalamus.

## Data analysis

### Orientation selectivity

In order to quantify the orientation selectivity (OS) of single cortical neurons, we calculate the preferred orientation (PO) and the orientation selectivity index (OSI). This information can be extracted from its respective tuning curve, *ν*(*θ*), representing the mean firing rate of a neuron in response to a stimulus of orientation *θ*. The method adopted here is to compute the orientation selectivity vector (OSV) as defined in circular statistics [26, 27]

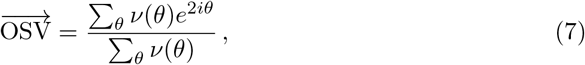

which is represented as a complex number. The PO corresponds to the phase (angle) of the OSV

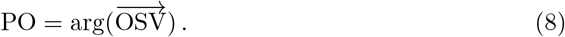

The OSI corresponds to the magnitude (length) of the OSV

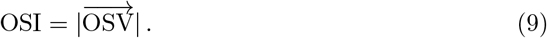

The OSI is frequently employed to describe the strength of orientation selectivity. A higher value of OSI means stronger orientation selectivity. A neuron has OSI = 1 when it selectively responds to one stimulus orientation and keeps silent for all other orientations. For an nonselective neuron responding equally to all orientations, we have OSI = 0.

In some experimental literature, an alternative measure of orientation selectivity is used

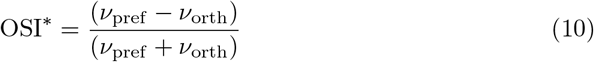

where *ν*_pref_ is the mean firing rate at the preferred orientation and *ν*_orth_ is the rate at its orthogonal orientation. Note that in previous theoretical work, it has been pointed out that, for a perfect cosine tuning curve, OSI^∗^ is twice as large as the OSI [28].

It is also necessary to figure out how measurement noise affects the level of OSI. To this end, we consider a stimulus of uniform luminosity (“zero contrast grating”) that is applied 12 times, each time with a different seed of the random number generator. As a result, the spike times of neurons are different although they are stimulated by exactly the same visual stimulus. The OSIs and POs are then extracted from the “tuning curves” built from these data. We performed 50 independent simulations to ensure robust results. The mean OSIs for both networks were around 0.047, and the POs of all 50 trials were uniformly distributed.

### Decoding neuronal responses

It is important that neuronal activity represents the stimulus orientation without a bias. To verify this, we decoded neuronal population activity in the V1 recurrent network to estimate the stimulus orientation using the population vector pCV for each stimulus orientation *θ*

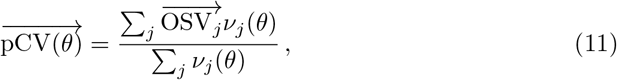

where 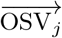 is the orientation selectivity vector of neuron *j* and *ν*_*j*_(*θ*) is the mean firing rate of neuron *j* at stimulus orientation *θ*. The angle of the population vector *θ*_pCV_ is the stimulus orientation decoded from the population activity, and the length of 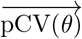 represents the confidence about the estimation (Fig 4). The value is between 0 and 1, where larger values signify higher confidence about the prediction.

To evaluate the deviation between the predicted orientation and the actual orientation of the stimulus, we calculated the circular difference between them (Fig 4A). The value of deviation is limited to the range [0^°^, 90^°^]. The deviation of 90^°^ means that the estimated orientation is perpendicular to the orientation of the stimulus and 0^°^ indicates a perfect agreement between them.

### Receptive field

Receptive fields are mapped using sparse noise stimuli and a reverse correlation procedure. For each stimulus image, an equal number of white and black spots are placed randomly in the visual field on the gray background matching the mean luminance. Around 20% of the visual field is filled with spots. In order to eliminate any boundary effects for neurons at the border of the visual field, the stimulus image is extended with gray background. The diameter of individual spots is 4^°^, roughly matching the size of the center of RGC receptive fields. 20 000 stimulus frames are used during simulation and each frame is presented for 33 ms. Receptive fields are then determined by a reverse correlation procedure, a commonly used method. All frames are averaged after weighting them by the neuronal response they evoked [29]. Note that no additional smoothing was applied to the figures of receptive fields.

## Results

To investigate the effects of cortico-thalamic (CT) feedback on the responses of cortical neurons and thalamic relay cells (TC), we performed numerical simulations of two networks: a feedforward thalamo-cortical network (without CT feedback) and thalamo-cortico-thalamic loop network (with CT feedback). Sinusoidal drifting gratings were used for visual stimulation (see Eq 3). The gratings were presented at 12 different orientations, covering the range between 0^°^ and 180^°^ in steps of 15^°^. The movement direction of the grating was always perpendicular to its orientation. In addition, the networks were also stimulated with sparse noise to determine the receptive fields of neurons. All numerical simulations were performed with NEST, a tool for the numerical simulation of spiking neural networks [30, 31].

Previous experimental and theoretical studies in mice lead to the conclusion that orientation selectivity of V1 neurons is essentially driven by inputs from non-selective dLGN relay cells [6, 18, 32, 33]. In our model, moving gratings induced activity in RGC cells with circular receptive fields, which oscillated with the temporal frequency of the grating. The mean (F0 component of the Fourier transform) and amplitude (F1 component) of temporal responses of single neuron were fixed, while their phases changed depending on stimulus orientation. Since in our model each TC cell receives input from exactly one RGC neuron, the same phenomena were also observed in the activity of individual TC cells. As a result, single TC neuron activity is not tuned to stimulus orientation. The receptive fields of V1 neurons were randomly positioned in the visual field, and each V1 neuron received input from multiple TC cells nearby. The compound thalamic input to individual V1 neurons was then a linear superposition of the activities of all presynaptic TC cells. Therefore, the compound signal is again a harmonic oscillation. The temporal mean of the compound signal does not depend on stimulus orientation, because the temporal mean of the activity of single TC cells is identical. However, its amplitude is generally tuned to the stimulus orientation depending on the phases of all presynaptic TC cells (see examples in S1 Fig.). These findings are fully compatible with experimental studies in mice [6]. Therefore, the orientation selectivity of thalamic inputs, 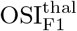 and 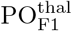, can be extracted from the orientation tuning curves of amplitudes.

### Orientation selectivity of V1 neurons is enhanced by CT feedback

To investigate the impact of CT feedback on the orientation selectivity of V1 neurons, we presented drifting sinusoidal gratings of 12 different orientations to our networks with and without CT feedback, respectively. Classically, orientation tuning is extracted from the mean firing rate of neuronal responses. We then extracted the orientation selectivity index (OSI) of all V1 neurons from their respective tuning curves, based on the circular variance (see Eq 9). A higher value of OSI indicates a more pronounced orientation selectivity. On the population level, the CT feedback led to higher mean OSI of V1 neurons (Fig 2 A). This effect was robust across all contrasts of the visual stimulation, but the amplification was stronger for intermediate contrasts. For stimuli with very low contrasts (< 0.1), however, the OSI approached the value obtained by measurement noise (see Methods).

**Fig 2.**
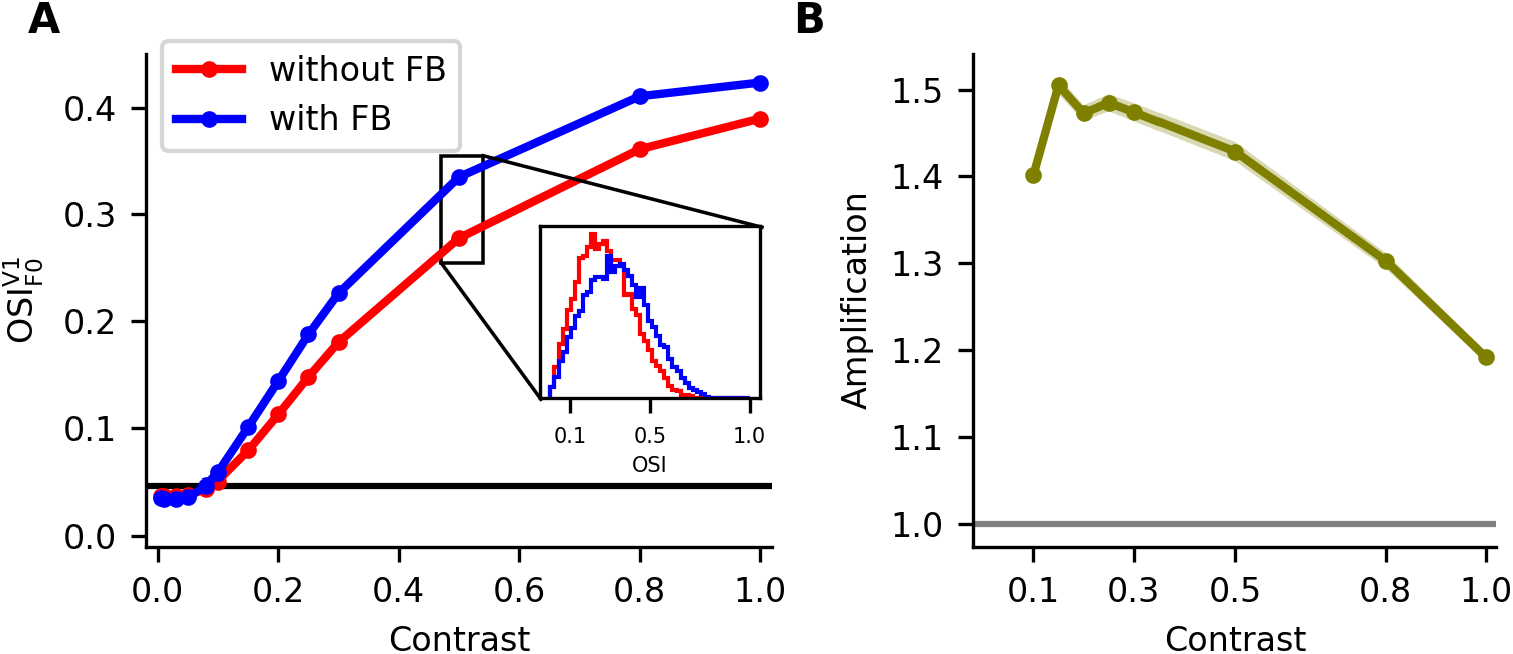
Amplification of OSI by CT feedback depends on contrast. A: The output OSIs of V1 neurons at different stimulus contrasts in networks with (blue) and without (red) CT feedback. The black line at around 0.047 indicates the OSI extracted from the presentation of zero contrast gratings (see Methods). The inset shows the OSI distribution of V1 neurons in networks at half contrast. B: The curve shows mean ± SEM of OSI amplifications of single V1 neurons. The amplification factor is calculated from the OSIs of single neurons by comparing the feedback with the feedforward network. Outliers for close to zero contrasts are deleted. The gray horizontal line at 1.0 indicates that the OSI is exactly the same in both networks. The solid lines and shaded areas with respective colors represent mean ± SEM.

Besides comparing the average OSI of the networks with and without CT feedback under different contrasts, we also investigated OSI amplification for individual V1 neurons. For a V1 neuron, the amplification factor (AF) of OSI indicates the relative increase of the OSI as a result of CT feedback. This number is 1 if there is no change, and it is larger than 1 if the OSI is larger in the network with feedback as compared to the network without feedback. We found that the single neuron AF of OSI was larger than 1 on average for all contrasts, although somewhat contrast dependent (Fig 2B). Remarkably, the AF was significantly larger for weaker contrasts (mean ≈ 1.5, median ≈ 1.26 for 0.15 contrast) as compared to full contrast (mean ≈ 1.2, median ≈ 1.08). To summarize, we find that the CT feedback had a strong effect on the strength of the orientation selectivity of V1 neurons, especially under conditions of weak stimulus contrasts. Note that the AF of neurons for the stimuli with very weak contrasts(< 0.1) is not shown here since the OSIs are too close to the value for zero contrast stimuli (see Methods).

### The correspondence of PO between input and output is better preserved in a network with feedback

Previous studies in different species have reported that the orientation selectivity of V1 neurons is essentially inherited from the orientation selectivity of their thalamic inputs [6, 32, 34]. We now asked whether the CT feedback contributes to this phenomenon. To this end, we first extract the orientation tuning curves of all V1 neurons and their thalamic inputs in the networks. We observed that the tuning curves of single V1 neurons are very similar to those of their thalamic inputs (Fig 3B) in both networks, while the similarities between input and output tuning curves are significantly higher in a network with CT feedback as compared to networks without CT feedback (*P* = 5.14 × 10^−68^, Wilcoxon test). To better understand the impact of CT feedback on orientation preference, we also extracted the preferred orientations (PO) of all neurons from their tuning curves. The result was that the PO of V1 neurons are more close to the PO of their input in the networks with CT feedback as compared to the feedforward network with CT feedback turned off (Fig 3C-D).

**Fig 3.**
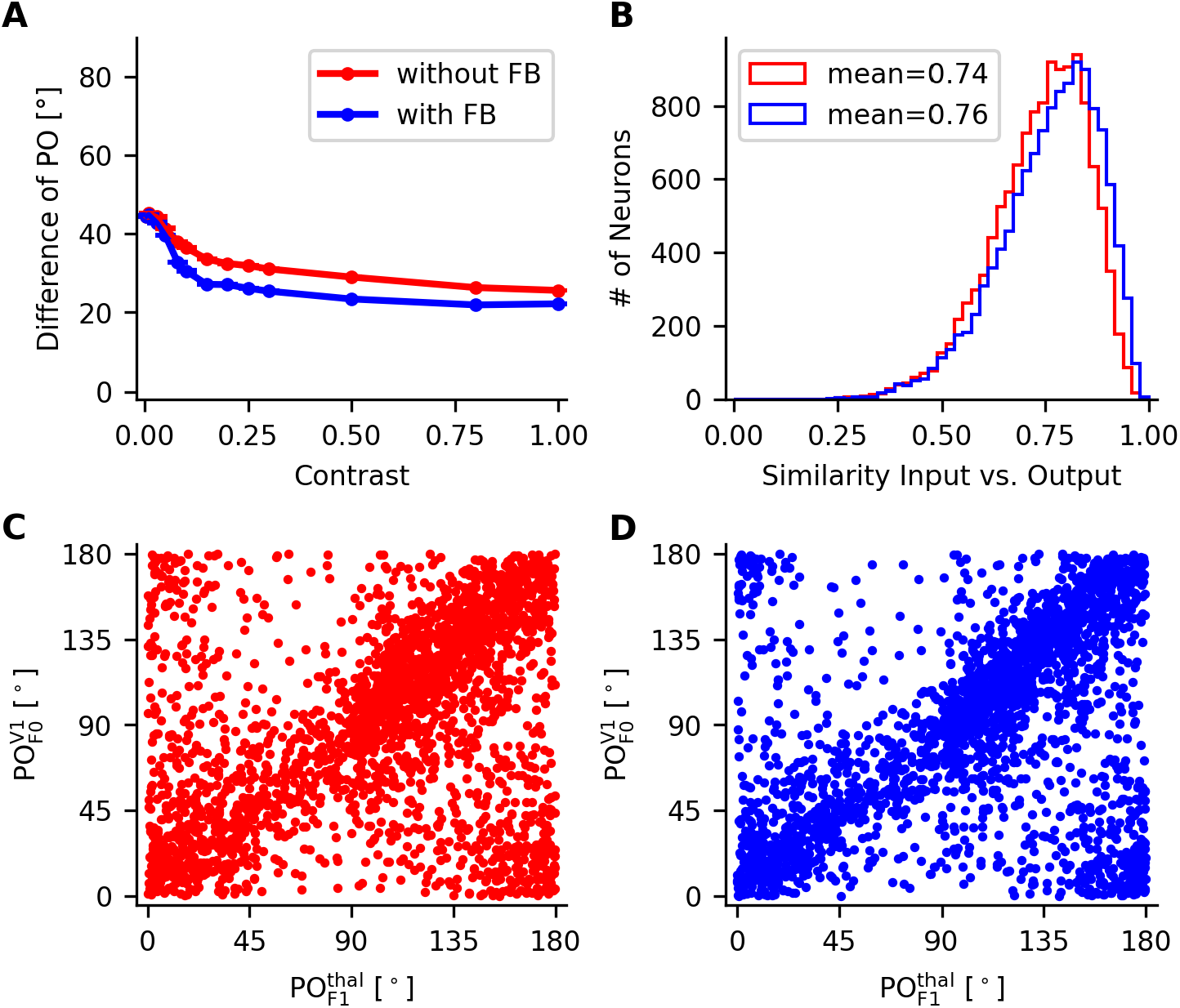
The PO is better transmitted from input to output for networks with CT feedback. A: Difference of input and output PO in networks with and without CT feedback for different stimulus contrasts. Dots indicate the population average. The respective SEM is very small. B: Distribution of similarities between input and output tuning curves of single V1 neurons in networks with and without CT feedback. C-D: Scatter plot of input and output PO in networks without (C, red) and with CT feedback (D, blue) at full contrast.

**Fig 4.**
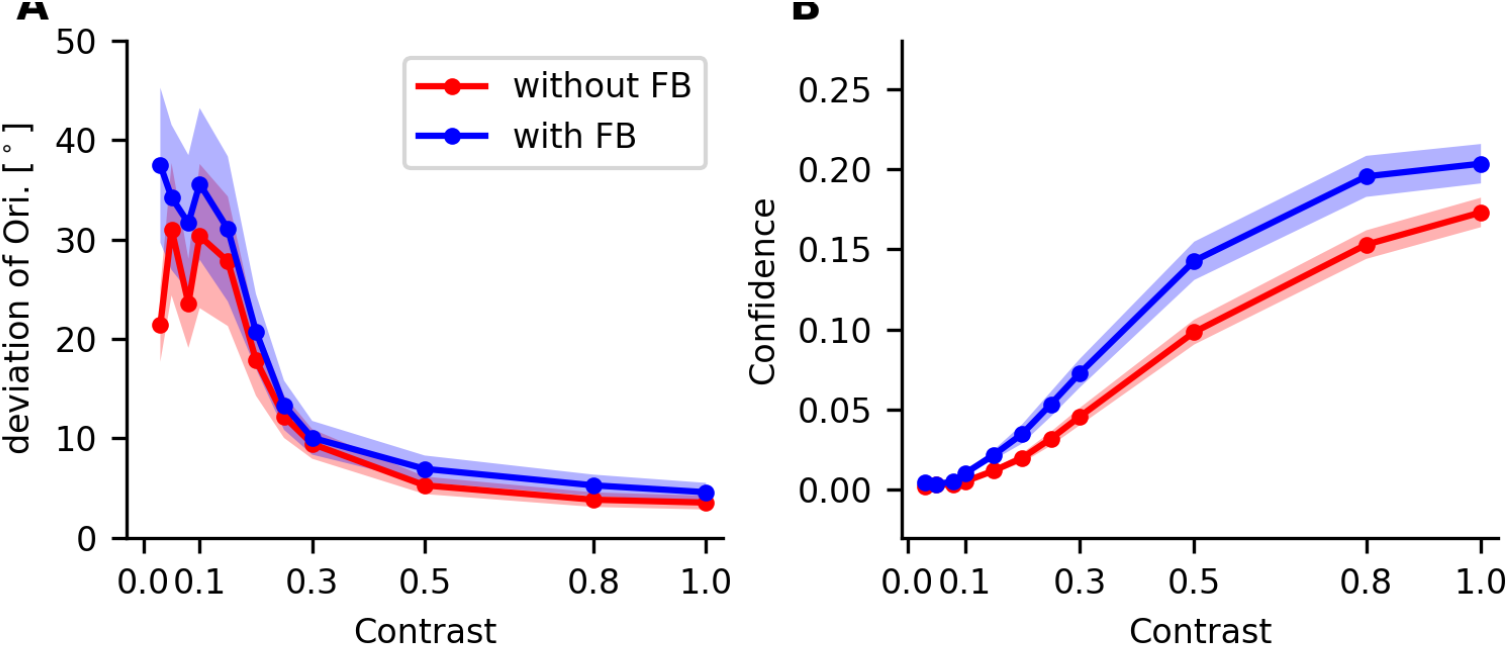
Decoding population activity. A: The deviation of estimated stimulus orientation and actual stimulus orientation in networks without (red) and with CT feedback (blue) are plotted against stimulus contrast. The smallest contrast considered is 0.03. B: Confidence values of population activity in both networks are plotted against the stimulus contrast. Solid lines with dots show the mean values and the respective shadows represent the SEM across 12 stimulus orientations.

To quantify the dependence of feedback effects on stimulus contrast, we calculated the circular differences of input and output PO of single V1 neurons at all contrasts ranging from 0 to 1. The circular difference between input and output PO is in the range between 0^°^ and 90^°^, where 0^°^ means that the output PO is perfectly aligned with its input PO, and 90^°^ means that they are orthogonal to each other. We found that the effect of CT feedback is robust (Fig 3A): The differences between input and output PO are generally smaller in the network with CT feedback across all contrasts, and this effect is even more pronounced for low contrasts. For very low contrasts (< 0.1), however, the orientation tuning is too weak to begin with to support a reliable estimation of the PO. The case of zero contrast is not shown in the figure as the stimulus then carries no information about orientation anyway. In summary, our analysis revealed that the relation between the input and output tuning is generally better preserved in a network with CT feedback.

### Neuronal population activity represents stimulus orientation

In previous sections, we have shown that the CT feedback not only enhances the orientation selectivity of neurons, but also improves the correspondence between the PO of input and output. To evaluate whether the CT feedback also affects the representation of the stimulus orientation in terms of neuronal population activity, we calculated the population vectors of networks with and without CT feedback. The population vector 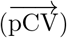 is obtained from the weighted sum of the orientation selectivity vectors 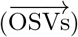 of individual neurons, normalized by the population rate (Eq 11). This also suggests a dependence on the OSI and PO of individual V1 neurons: The population PO is the orientation of 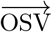 and the population OSI is the length of 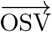. Therefore, we hypothesized that the network with CT feedback should represent the stimulus orientation better. To test our hypothesis, we extracted two parameters from the population vector: the angle of the vector is the stimulus orientation represented by the population activity and its length is the confidence assigned to the representation. Again, we extracted these variables for all stimulus contrasts in both networks.

To compare the represented orientation with the actual orientation of a stimulus, we calculated the circular difference between them. We found that the deviation is relative large for very weak contrasts and decreased rapidly for stronger contrasts in both types of networks (Fig 4 A). This means that neuronal population activity of V1 neurons can represent the stimulus orientations provided the stimulus contrast is larger than 0.15. Although the estimated orientation deviated slightly more from the actual stimulus orientations in the networks with feedback as compared to the pure feedforward network, the confidence was significantly increased by feedback, for all contrasts (Fig 4B). Not surprisingly, the confidence values are generally increasing for higher contrasts. In summary, we find that the neuronal population activity in a network with feedback can represent the stimulus orientation with higher confidence than in the feedforward network.

### Signal-to-noise ratio of thalamic input is enhanced by CT feedback

We showed that the orientation selectivity of V1 neurons is enhanced by CT feedback, and that the tuning of cortical output is strongly correlated to the tuning of thalamic inputs. This raises the question whether thalamic inputs converging to individual V1 neurons are changed by the CT feedback. Since the compound thalamic input to V1 neurons was oscillating with the temporal frequency of the drifting grating, we quantified the thalamic input by its mean (the F0 component) and its amplitude at the fundamental frequency (the F1 component). The signal-to-noise ratio 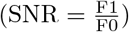 quantifies the relative strength of the temporal modulation. To evaluate the impact of CT feedback on thalamic inputs, we again calculated the amplification factors of these three parameters at different contrasts, which quantified the ratio of them in the network with feedback as compared to the network without feedback. The distributions of AFs at full contrast revealed different modulations in F0 and F1 components: The F0 components were suppressed (Fig 5D) while the F1 components were mostly enhanced in the network with feedback (Fig 5C) and therefore induced a higher SNR (Fig 5B). These effects on the amplification factors were robust across stimulus contrasts. However, the contrast dependence of the CT feedback was different (Fig 5 A): While the F1 and SNR are stronger amplified at lower contrasts, the reduction of the F0 component was almost the same for all contrasts.

**Fig 5.**
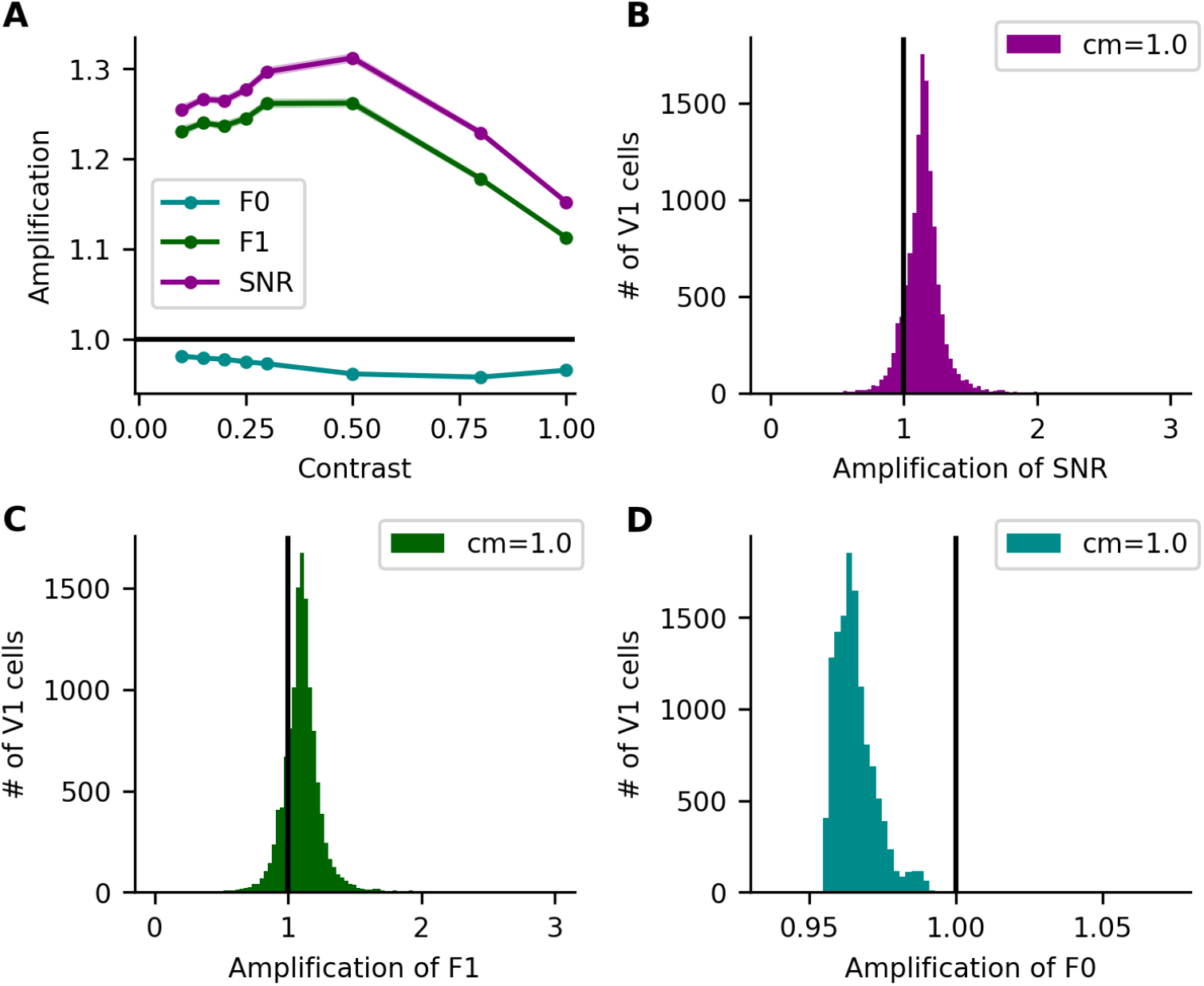
Contrast-dependent effects of CT feedback on thalamic inputs. A: The amplification factors of F0, F1 and SNR are contrast dependent. Shown are mean ± SEM (dots and shadows, respectively) of all three parameters for all thalamic inputs against stimulus contrasts. B-D: Distribution of amplification factors in the SNR (B), F1 component (C) and F0 component (D) at full contrasts. The horizontal (A) and vertical (B-D) black lines at 1.0 indicate no change as a result of CT feedback.

In the previous section, we verified that the F0 component of the thalamic input essentially conveys no information about stimulus orientation (see examples in S1 Fig.).

Therefore, our analyses indicate that CT feedback enhances the SNR of thalamic inputs by amplifying the component tuned to orientation (F1) and suppressing the untuned part (F0). The amplification of the SNR of the inputs by CT feedback are more pronounced for weaker contrasts, consistent with their V1 outputs (Fig 2).

### The SNR of single TC cells is increased by CT feedback

Compound thalamic input in our model is a linear superposition of all inputs from presynaptic TC neurons. Therefore, we further investigated the effects of CT feedback on the responses of single TC neurons. We first extracted the F0 and F1 components from the temporal responses of TC cells and calculated the SNR. The AF of these three parameters were then calculated. At full contrast, the distributions of the AF of TC cells differed from the AF of compound thalamic inputs to V1 neurons: Most of the F0 (Fig 6D) and F1 components (Fig 6C) were smaller in the feedback network, while the SNR was mostly larger in networks with feedback since the F1 components of single TC cells were less attenuated than the corresponding F0 (Fig 6B). Consequently, the CT feedback effects on these parameters were found to be contrast dependent (Fig 6A). The AF of F1 and SNR were larger for lower contrasts compared to the case of full contrast. We found that although the F1 components were slightly suppressed in the feedback network at full contrast, they were amplified at contrasts below 0.8 and reached the peak at 0.5. The reduction of F0 components was stronger for increasing contrasts and became again weaker at full contrast. As a result, the amplification effect of SNR was the strongest at the intermediate stimulus contrasts.

**Fig 6.**
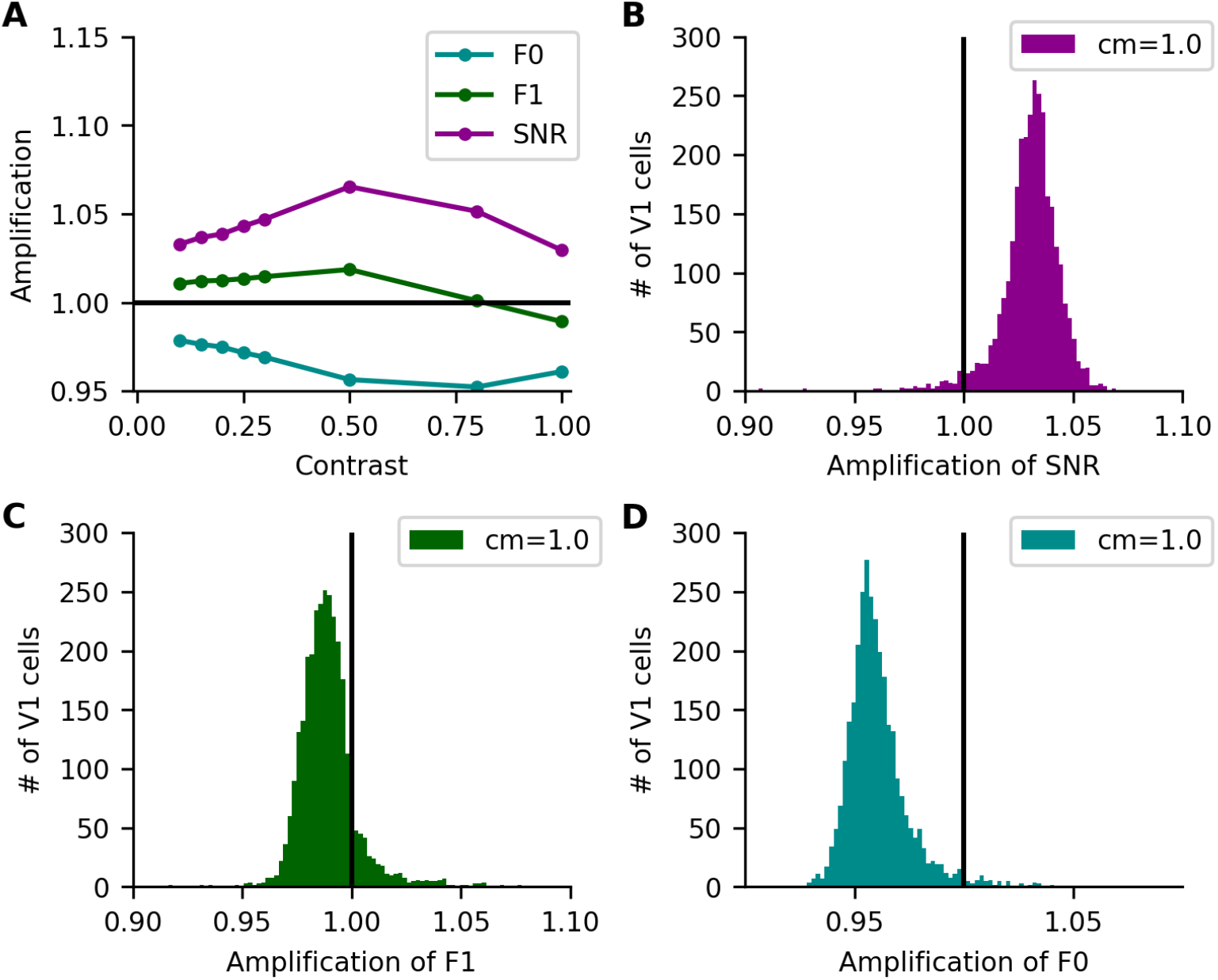
Effects of CT feedback on single TC cells. A: Contrast-dependence of the amplification factors of F0, F1 and SNR. The dots and shadows indicate mean ± SEM of these parameters. B-D: The distributions of the amplification factors of SNR (B), F1 (C) and F0 (D) in individual TC neurons. The horizontal and vertical lines are the same as in Fig 5.

The AF of F0 components of single TC neurons and the compound thalamic inputs were distributed in the same range, while the AF of F1 and SNR of TC neurons were in a narrower range compared to thalamic inputs. This can be explained by the linear superposition of TC response curves. The F0 component of the compound input was only determined by the F0 of individual responses, while the F1 of the compound thalamic signal was affected not only by the F1 of individual neurons, but also the phases of the their response curves. This indicates that although the modulations of CT feedback on the responses of TC cells might be weak, the proper summation of presynaptic responses, however, can induce a significant change.

### Cortico-thalamic feedback enhances the receptive fields of TC neurons

A series of experimental studies revealed that CT feedback modifies the receptive fields of dLGN neurons [9, 15, 17, 35]. In our model, CT feedback indeed modulates the responses of TC cells in the dLGN. Therefore, it seemed promising to study the role of CT feedback on the receptive fields of TC neurons. We stimulated networks with and without feedback with sparse noise stimuli (see S2 Fig.) and then characterized the receptive fields of TC neurons by calculating the spike-triggered averages (STA). The RF of a sample OFF center cell showed clear center-surround structure under both conditions (Fig 7A). Although they seemed to have roughly similar shapes, the RF center is indeed stronger in the network with feedback. In order to visualize the change of the RF as a result of feedback, we extracted their peak values and plotted the distributions of them (Fig 7B). Across the population, the RF peak amplitudes was mostly smaller in the network without feedback. The distributions of the peak values of the differences between the RFs in both networks (Fig 7B, inset) indicated clearly that the center of the RFs of TC neurons were stronger in the network with feedback. To further investigate the effect of CT feedback on the RFs of V1 neurons, we extracted them and their respective thalamic inputs in both scenarios (Fig 7C). The RFs of V1 neurons were strongly correlated with the RFs of their thalamic inputs. However, the differences of the RFs in networks without and with feedback were not significant. We conclude that CT feedback indeed enhanced the receptive fields of TC cells, while the RF of V1 neurons were not strongly affected.

**Fig 7.**
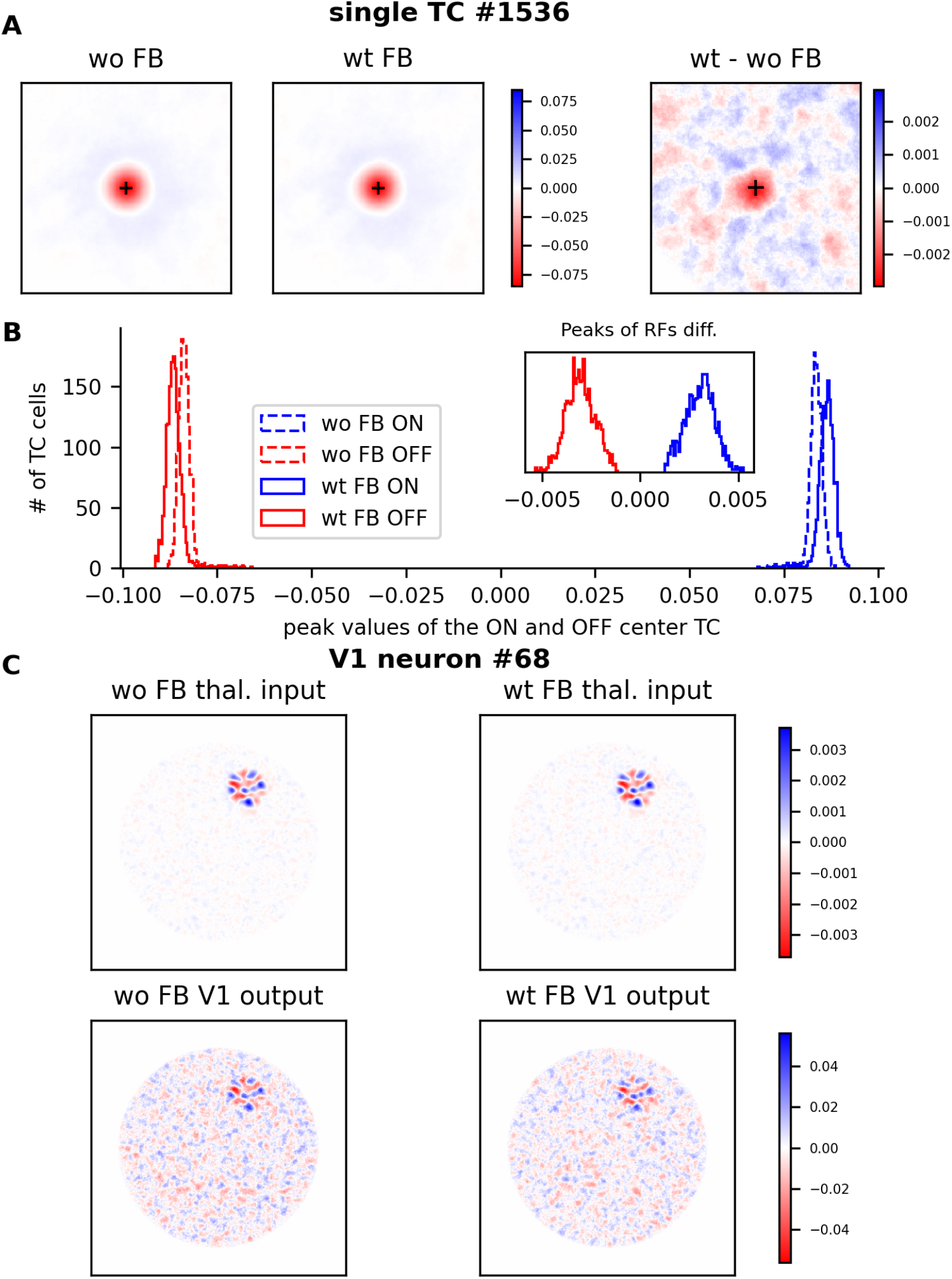
Receptive field structures of sample neurons. A: The RFs of an OFF center TC cell (id #1536) in a network with (wt, middle) and without (wo, left) CT feedback. The right panel shows the difference between the two RFs. B: Distribution of the peak values of the RFs of ON and OFF TC cells in feedforward and feedback networks. The inset histogram shows the distribution of the peak of RF differences. Blue and red colors indicate ON and OFF RFs, respectively. C: The RFs of a sample V1 cell (id #68) and its thalamic inputs without and with feedback.

### The geometry of CT feedback affects the OS of V1 neurons

In the feedback pathway, the cortico-thalamic axons project to TC neurons (direct excitation) as well as GABAergic interneurons and TRN neurons, which further innervate TC neurons (indirect inhibition). The reciprocal connections between TC and TRN neurons are mostly open-loop connections, forming the substrate of lateral inhibition [19, 36]. Although TRN neurons can also be excited by TC inputs, they are mostly driven by the CT excitation [37]. This general configuration suggests that the spatial arrangement of CT feedback might have an important role in modulating thalamo-cortical transmission. To illustrate this hypothesis, we applied two somewhat extreme connection rules for the direct and indirect feedback pathways. The “nearest” rule is based on the distances between receptive fields, and “random” projections cover the whole visual field and choose their targets with uniform probability. As a result, CT feedback connections end up in four different combinations: nearest direct excitation and nearest indirect inhibition (NE+NI), nearest direct excitation and random indirect inhibition (NE+RI), random direct excitation and nearest indirect inhibition (RE+RI) and random direct excitation and random indirect inhibition (RE+NI). We then extracted the tuning curves of V1 neurons from simulations of the four specific configurations to calculate the OSI and PO (Fig 8).

**Fig 8.**
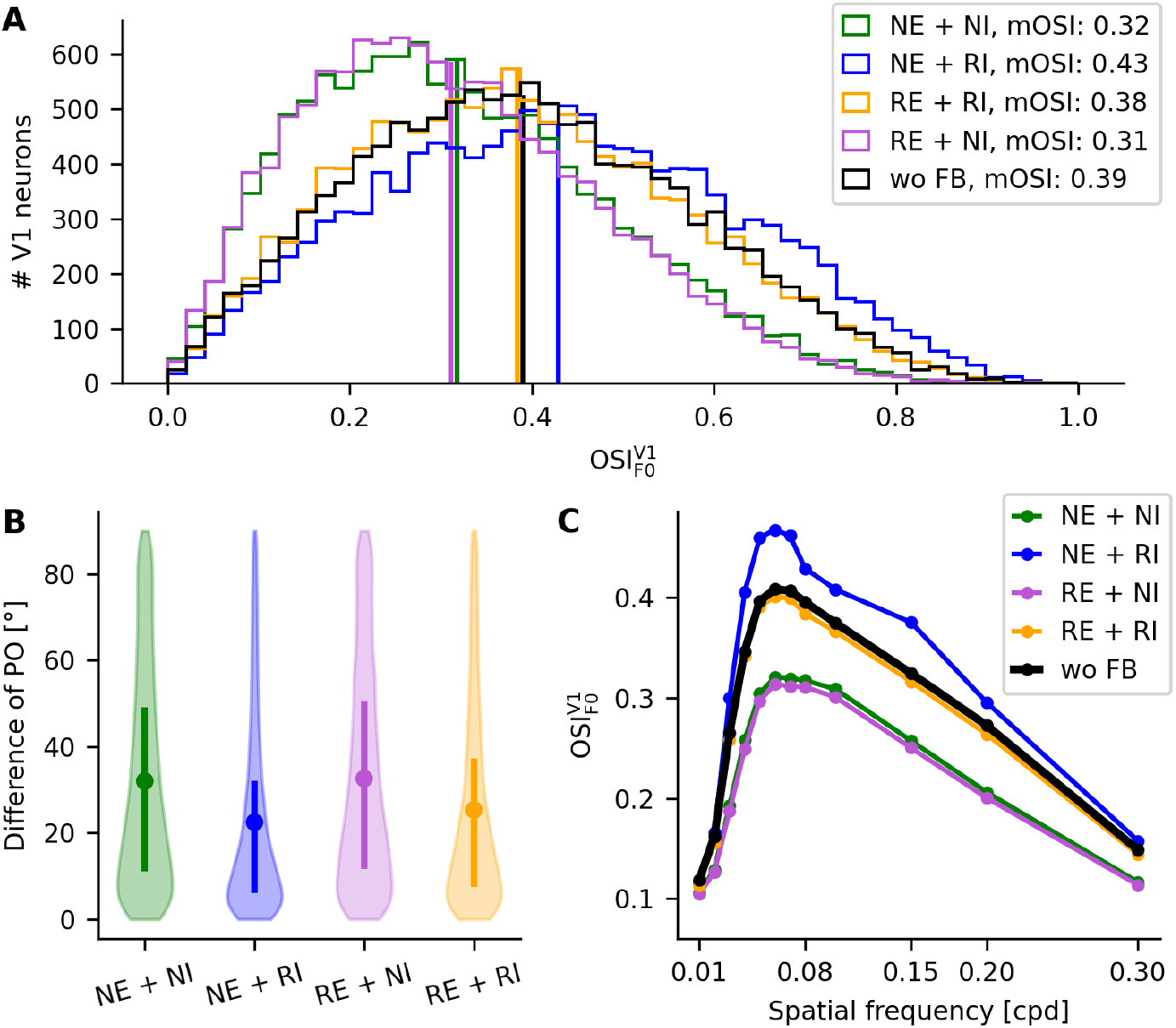
The geometry of feedback projections has a strong effect on the OSI of V1 neurons. A: Histogram of the OSI values of V1 neurons in networks with different geometries of the CT feedback. Vertical lines indicate the mean OSI of the respective distribution. B: Distribution of the difference of input and output PO of V1 neurons with different connection rules. The bars indicate the percentile range between 25% and 75% of the data, respectively. C: Dependency of the OSI on the spatial frequency in the feedback network with different connection rules, and the network without CT feedback. The dotted lines and shadows represent the respective mean ± SEM. NE: nearest direct excitation; NI: nearest indirect inhibition; RE: random direct excitation; RI: random indirect inhibition. Black color represents the results for the purely feedforward network.

As expected, the OSI indeed depended on the spatial configuration of CT feedback (Fig 8A). For the feedback with NE+RI, the neurons have stronger orientation selectivity than in the other settings. The distribution of OSI in a feedback network with RE+NI was very similar to the network without feedback. The OSI for the two regimes with nearest indirect inhibition (NI) have even reduced OS compared to networks without feedback. If feedback inhibition obeys the nearest rule, it conveys orientation-tuned information, similar to excitation. Therefore, it attenuated the transmission of orientation tuning. In addition, the input PO of V1 neurons were more distorted in the presence of feedback with nearest indirect inhibition (Fig 8B, green and purple).

Beyond studying the impact of feedback geometry on orientation selectivity for a fixed spatial frequency, we verified that the overall effects were robust across a large range of spatial frequencies (Fig 8C). In contrast, at small spatial frequencies with poor orientation selectivity below 0.02 cycles per degree (cpd), the facilitation of CT feedback in the configuration NE+RI was less pronounced.

### The effect of the strength of CT feedback on the modulation of V1 neuronal activity

Our analyses revealed that CT feedback enhanced the SNR of the thalamic inputs, which amplified the orientation selectivity of V1 neurons. The effects were robust for all stimulus contrasts but most pronounced at intermediate contrasts. In our network configuration, each CT neuron in V1 projects to a fixed number of neurons in the postsynaptic populations. Therefore, the total strength of CT feedback is proportional to the number of CT neurons in V1 (N_fb_). This raises the question how exactly the amount of CT feedback exerts its effects. To answer this question, we simulated different networks with a wide range of N_fb_ between 0 and 7 000 for different contrasts. If N_fb_ = 0, the CT feedback is effectively turned off and it corresponds to a feedforward network.

First, we calculated the population vector to test whether the neuronal population activity can represent the orientation of the stimulus correctly under different conditions. For full contrast, when we compared the orientation of the population vector to the actual stimulus orientation, we found that deviations between them were relatively small and increased slowly with N_fb_ until 5 000 and steeply for even larger values (Fig 9A). For weaker contrasts, the deviations remained similar while the kink shifted to smaller values of N_fb_ (Fig 9B-D): the turning point was N_fb_ = 5 000 for full contrast and shifted to 3 500 at half contrast. However, the deviations at 0.1 contrast are large for all values of N_fb_, including the no-feedback network N_fb_ = 0 (Fig 9E). Under this condition, the orientation bias in the input was too weak to be transformed into the output. Similar to the case of zero contrast stimuli, extracting a PO from the tuning curves becomes impossible.

**Fig 9.**
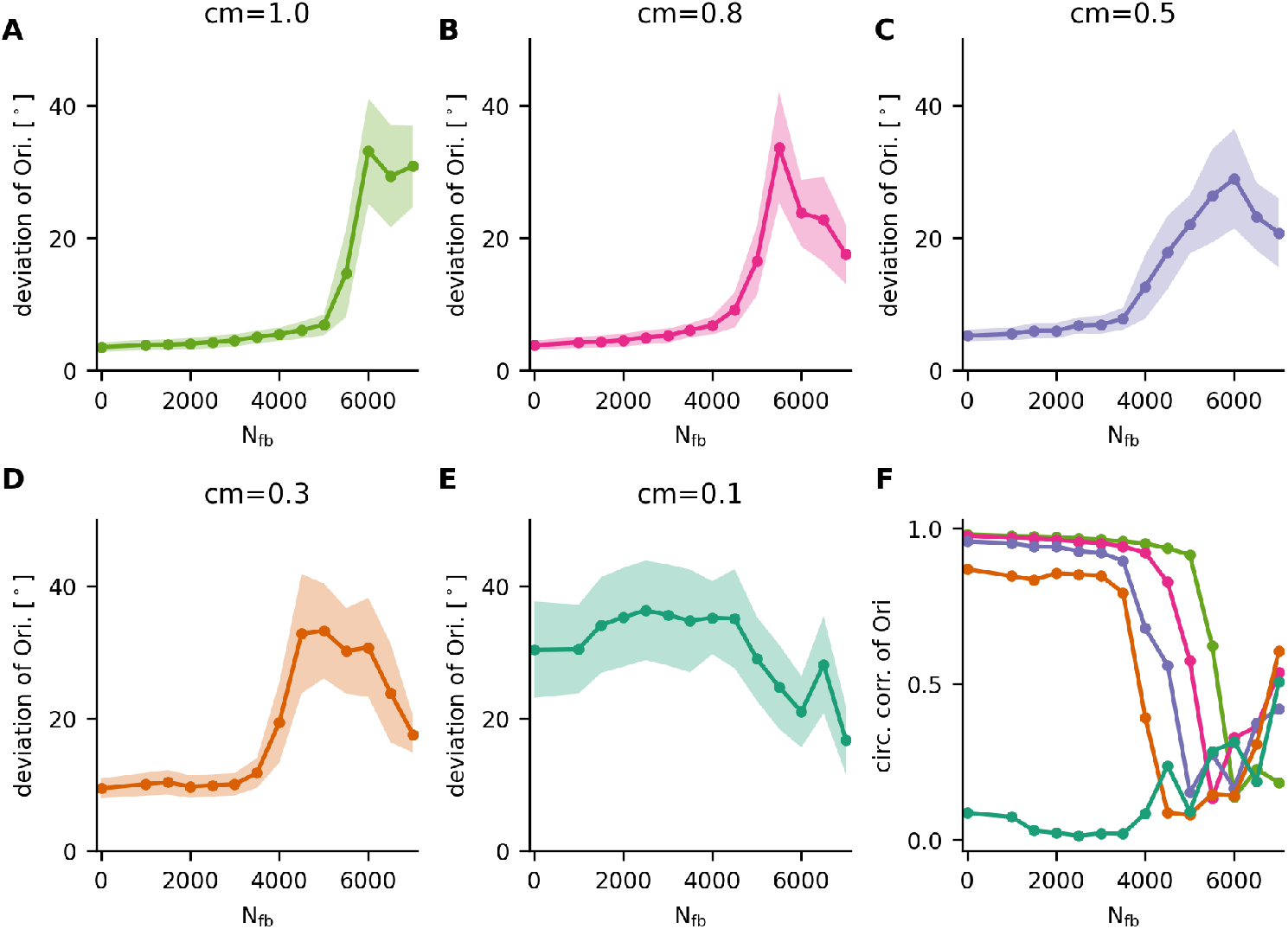
The representation of stimulus orientation depends on the strength of feedback and stimulus contrast. A-E: Deviations between the actual stimulus orientation and the orientation represented by population activity in the network, for different values of N_fb_ and for different stimulus contrasts. Dots and shadows show the respective mean ± STD. F: Circular correlation between the represented orientation and the actual stimulus orientation. The colored curves indicate the same contrast as displayed in panel A-E. Larger values of the circular correlation indicate better representation of the actual stimulus orientation by the population.

Finally, we investigated the firing rates of V1 neurons in the different simulation scenarios. We found that the mean and the standard deviation of firing rates depended on N_fb_, very much like the deviations between represented and actual stimulus orientations (Fig 10A-B): Both parameters first increased slowly with N_fb_ and then grew abruptly for larger values. For weaker contrasts, the turning point moved to smaller values of N_fb_. For extreme values of N_fb_, most V1 neurons were firing at relative low frequencies (≈ 1 Hz), but some at extremely high rates (≈320 Hz). This suggested that under these conditions, a small group of V1 neurons made a huge contribution to the population activity, which cannot generate a faithful representation of the stimulus any more. To assess the orientation preference of the network, we extracted the the distribution of POs of all V1 neurons using a histogram with bins of width 1^°^. Ideally, the POs are uniformly distributed (maximum entropy, *H*_max_ = log(180) ≈ 5.19), and the population has no bias in its orientation preference. Fig 10C shows that the entropy of the PO distribution was close to the maximum for smaller values of N_fb_, but it decreased sharply for higher values N_fb_ beyond some threshold. This indicated that for very large values of N_fb_, the POs of V1 neurons are non-uniformly distributed, which induces orientation-selective responses of the population that are unrelated to the stimulus. In summary, the strength of CT feedback does affect the activity of V1 neurons, which induces a strong bias in the orientation selectivity of the population. All these effects are contrast dependent.

**Fig 10.**
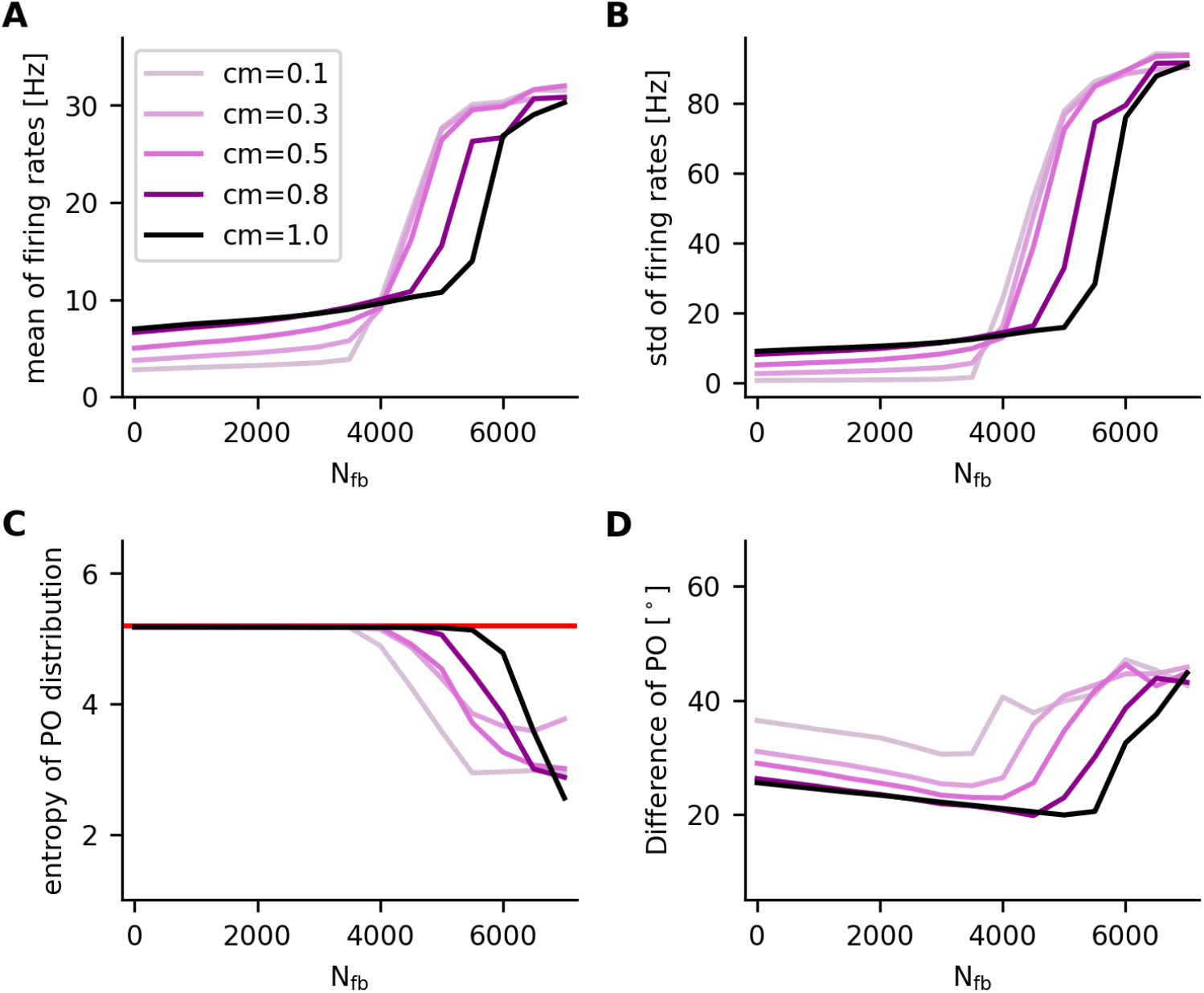
A,B: Mean (A) and standard deviation (STD, B) of network firing rates over all neurons in the networks for different feedback N_fb_ and different stimulus contrasts. C: Entropy of the distribution of POs of all V1 neurons. The red horizontal line at log(180) ≈ 5.19 indicates the entropy of a uniform distribution with 180 bins of width 1^°^. D: The differences between the input and output PO for different configurations of the network.

## Discussion

The aim of this paper is to elucidate the functional role of CT feedback in visual processing. We focused on the effect of feedback for selective responses of cortical neurons and their underlying mechanisms. To this end, we have developed a spiking network model of the thalamo-cortico-thalamic loop circuits based on our previous work [18], in which orientation selective responses of V1 neurons emerge in the feedforward pathway. Using sinusoidal moving gratings at different orientations as a stimulus to the network, we observed that the CT feedback enhances the orientation selectivity (OS) of V1 neurons without distorting their thalamic inputs. As a result, the correspondence between input and output PO is better preserved, and the population activity of V1 neurons represents the stimulus orientation with higher confidence. Importantly, these effects are more pronounced for lower stimulus contrasts.

Since the preferred orientation of cortical outputs is highly correlated to the PO of their thalamic inputs, our analysis of the activity of TC cells revealed a potential mechanism for the impact of feedback. The SNRs of the temporal responses of TC cells are generally increased by CT feedback, which is consistent with experimental findings [38], and which results in a higher SNR of the compound thalamic inputs to V1 neurons. Furthermore, we have shown that CT feedback enhances responses of TC cells to a stimulation of the center of their RF.

In addition, our simulation results indicate that these enhancing effects on the coding of CT neurons are highly dependent on the spatial profile and the number of feedback projections. The connections of direct excitation and indirect inhibition need to be arranged properly to amplify the orientation preference of V1 neurons, as will be discussed in more detail later.

In most experimental studies, the visual cortex was silenced to examine the impact of CT feedback on neurons in subcortical regions, such as dLGN and TRN [14, 17, 38–42]. However, there are still very few experiments that directly address the effect of feedback on the tuning of V1 neurons, which receive their visual input mainly from dLGN neurons. In fact, those feedback inputs are often blocked in experiments. The computational model, in contrast, enables us to record and analyze neuronal activity across all populations, addressing the sensory processing directly and thus elucidating its underlying mechanisms.

### Firing mode of TC cells and CT feedback effects

The relay cells in the dLGN exhibit two different firing modes, burst and tonic, which are thought to have a pronounced impact on the transmission of visual information. The tonic mode is associated with undistorted linear transmission of visual inputs, while burst firing can induce a nonlinear transformation of the signals with increased efficacy [43–46]. In addition to stimulus-driven bursts, thalamic bursting is mainly triggered by the de-inactivation of low-threshold T-type calcium channels. [43, 47].

In this study, we employed standard current-based leaky integrate-and-fire (LIF) neuron models. We did not record prominent burst spikes in TC neurons, as this simple model lacks calcium channels which precludes the generation of such bursting activity. However, our analyses showed that the CT feedback primarily enhanced the signal-to-noise ratio (SNR) of the temporal responses of TC neurons, which amplifies visual transmission, consistent with experimental results [38]. In our model, this augmentation is one of the mechanisms underlying the stronger SNR of the compound thalamic signal. Another mechanism is the synchronization of temporal responses of presynaptic TC neurons.

Some studies have reported that the CT feedback has no significant impact on the proportion of burst spikes [38, 48], while others have observed that the CT feedback facilitates the transition of the firing mode from burst to tonic [12, 14, 49]. These discrepancies in conclusions might be a result of variations in experimental and analytical methods. When the firing mode of TC neurons undergoes a subtle shift from burst to tonic, it allows a higher dynamic range for the transmission of visual information. The impact of the firing mode change on CT feedback is indeed bidirectional: It shifts the response phases of TC neurons, which in turn affects the response synchrony of presynaptic TC neurons.

### Geometry of CT feedback connections and receptive fields

Pyramidal cells in the cortex project to TC cells in dLGN through direct excitatory and indirect inhibitory pathways. The geometry of these two projections may be critical for their function, as it has a strong impact on the balance between feedback excitation and inhibition.

In our study, we explored the effects of four distinct hypothetical geometries of cortico-thalamic (CT) feedback on the orientation selectivity of V1 neurons and compared them to a purely feedforward network without any feedback. These four spatial configurations result from two extreme connectivity rules to each feedback pathway. The “nearest rule” assumes that CT neurons project to neurons whose receptive fields (RFs) are closest, whereas the “random rule” prescribes that CT neurons select targets from the entire visual field with uniform probability. Measuring the orientation preference of V1 neurons under all these conditions, we found that the exact spatial arrangement of CT feedback indeed has an impact on the selective responses of V1 neurons. Specifically, when the direct excitatory projections follow the “nearest” rule while the indirect inhibitory inputs originate randomly from the visual field, the CT feedback induces a pronounced amplification of orientation selectivity in V1 neurons. These findings suggest that a combination of narrow direct excitation and unfocused indirect inhibition constitutes a promising strategy for enhancing the efficiency of visual transmission. This spatial motif indeed aligns well with experimental findings: Direct feedback excitation is more localized, while indirect feedback inhibition via TRN neurons exhibits a broad spatial dispersion [3, 17, 50].

The spatial structure of receptive fields depends essentially on synaptic inputs. While the RFs of TC neurons closely resemble the RFs of RGCs, they are modulated by different feedback inputs from cortex. Therefore, the interplay between direct excitation and indirect inhibition may play an important role in shaping the RF of TC cells as well. Using sparse noise stimuli and reverse correlation techniques, we found that the centers of the TC RFs are amplified by the CT feedback under the condition NE+RI, i.e. nearest direct excitation and unfocused indirect inhibition. Experiments indeed generally report that the CT feedback amplifies surround suppression [17, 51, 52]. A direct quantitative comparison of our simulations with experimental findings is problematic because different types of visual stimuli and analysis methods are used in the different experiments. More importantly, inhibitory inputs converging to TC neurons cover the entire visual field under the NE+RI condition, resulting in wider and weaker inhibition. This potentially attenuates the effect of cortical feedback on surround suppression.

### Cortical network model

In rodent brains, the thalamo-cortical axons in dLGN relay cells predominantly target neurons in layer 4 of the primary visual cortex, while the majority of cortico-thalamic feedback to TC cells originates from layer 6 of V1. It has been shown in several studies that neurons in L4 and L6 exhibit distinct activities and connectivity [53–56]. For instance, the L6 neurons fire at lower rates and exhibit weaker orientation selectivity than L4 neurons. These properties might contribute to the balance of the feedback excitation and inhibition and the orientation selectivity of relay cells in dLGN.

In our model, a number of simplifications were made with regard to anatomy. In particular, we employed a simple recurrent network without layers and layer-specific projections to model the V1 network [22]. The cortico-thalamic neurons in our model are a subset of excitatory neurons chosen randomly from the V1 recurrent network. As a result, the activities of CT neurons and other excitatory neurons in the recurrent network are the same. Despite its simplicity, our model represents a wide range of properties such as linear and nonlinear transformation of the visual information, and it allows to study how orientation selectivity emerges, and which factors modulate it.

## Supporting information

**S1 Fig.**
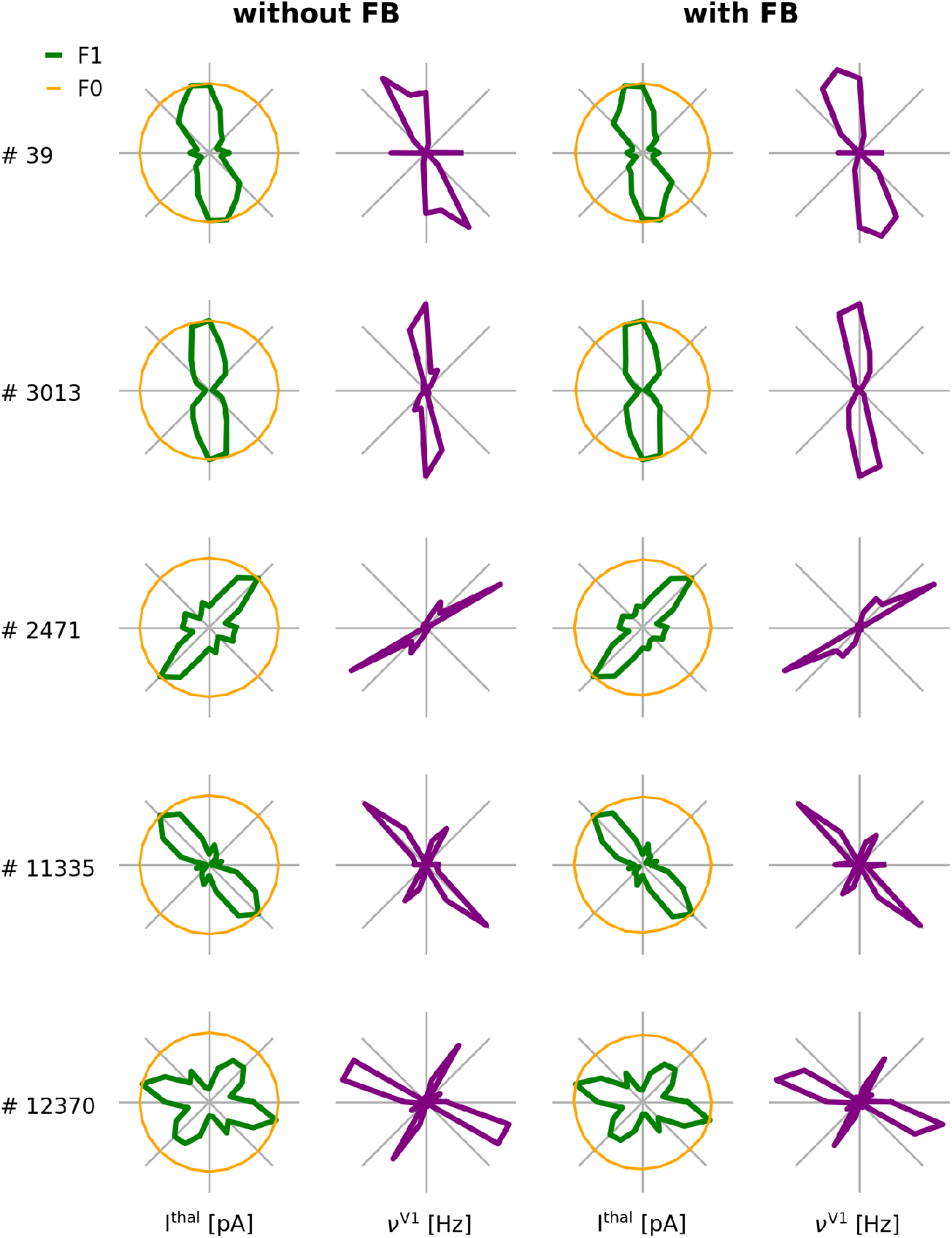
Tuning curves of V1 neurons and their thalamic inputs in networks with and without CT feedback. Examples of matching input (1st and 3rd columns) and output (2nd and 4th columns) orientation tuning curves in networks with and without CT feedback in polar representation (360^°^). Orange and green curves represent the normalized F0 and F1 components of the thalamic input currents, and purple curves show the tuning curves of the output mean firing rates of sample V1 neurons.

**S2 Fig.**
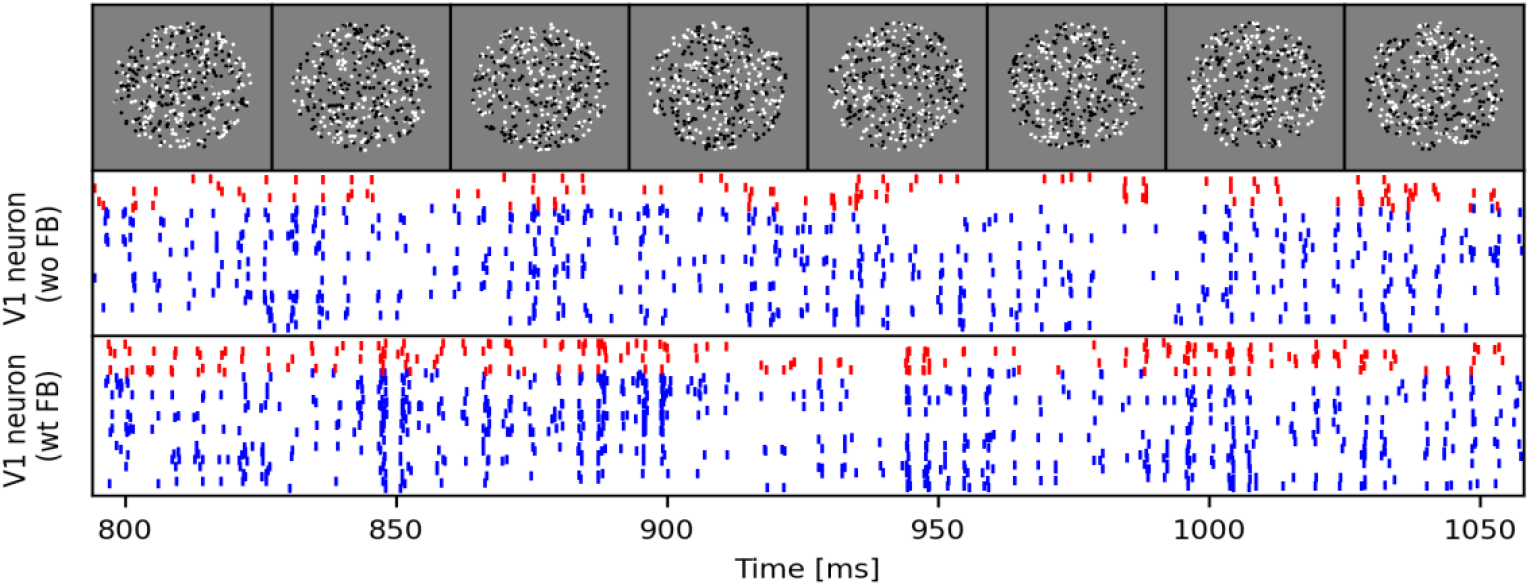
Sparse noise stimuli used to map receptive fields. The spike trains of sample V1 neuron responses to the shown stimuli (top) in the feedforward and feedback network, respectively. Blue and red spikes correspond to excitatory and inhibitory neurons, respectively.

## Acknowledgments

This work was funded by the University of Freiburg. The HPC facilities used for this work are funded by the state of Baden-Württemberg through bwHPC and DFG grant INST 39/963-1 FUGG. The funders had no role in study design, data collection and analysis, decision to publish, or preparation of the manuscript.

## References

1. Sillito AM, Jones HE. Corticothalamic interactions in the transfer of visual information. Philosophical Transactions of the Royal Society of London Series B: Biological Sciences. 2002;357(1428):1739–1752.

2. Harris KD, Mrsic-Flogel TD. Cortical connectivity and sensory coding. Nature. 2013;503(7474):51–58.

3. Briggs F. Role of feedback connections in central visual processing. Annual review of vision science. 2020;6:313–334.

4. Hubel DH, Wiesel TN. Receptive fields, binocular interaction and functional architecture in the cat’s visual cortex. The Journal of physiology. 1962;160(1):106.

5. Ferster D, Chung S, Wheat H. Orientation selectivity of thalamic input to simple cells of cat visual cortex. Nature. 1996;380(6571):249–252.

6. Lien AD, Scanziani M. Tuned thalamic excitation is amplified by visual cortical circuits. Nature neuroscience. 2013;16(9):1315–1323.

7. Sherman SM, Guillery R. On the actions that one nerve cell can have on another: distinguishing “drivers” from “modulators”. Proceedings of the National Academy of Sciences. 1998;95(12):7121–7126.

8. Squire RF, Noudoost B, Schafer RJ, Moore T. Prefrontal contributions to visual selective attention. Annual review of neuroscience. 2013;36:451–466.

9. Briggs F, Usrey WM. Emerging views of corticothalamic function. Current opinion in neurobiology. 2008;18(4):403–407.

10. Alitto HJ, Usrey WM. Corticothalamic feedback and sensory processing. Current opinion in neurobiology. 2003;13(4):440–445.

11. Cudeiro J, Sillito AM. Looking back: corticothalamic feedback and early visual processing. Trends in neurosciences. 2006;29(6):298–306.

12. Wang W, Jones HE, Andolina IM, Salt TE, Sillito AM. Functional alignment of feedback effects from visual cortex to thalamus. Nature neuroscience. 2006;9(10):1330–1336.

13. Augustinaite S, Kuhn B, Helm PJ, Heggelund P. NMDA spike/plateau potentials in dendrites of thalamocortical neurons. Journal of Neuroscience. 2014;34(33):10892–10905.

14. Spacek MA, Crombie D, Bauer Y, Born G, Liu X, Katzner S, et al. Robust effects of corticothalamic feedback and behavioral state on movie responses in mouse dLGN. Elife. 2022;11:e70469.

15. Murphy P, Sillito A. Corticofugal feedback influences the generation of length tuning in the visual pathway. Nature. 1987;329(6141):727–729.

16. Hasse JM, Briggs F. Corticogeniculate feedback sharpens the temporal precision and spatial resolution of visual signals in the ferret. Proceedings of the National Academy of Sciences. 2017;114(30):E6222–E6230.

17. Born G, Schneider-Soupiadis FA, Erisken S, Vaiceliunaite A, Lao CL, Mobarhan MH, et al. Corticothalamic feedback sculpts visual spatial integration in mouse thalamus. Nature neuroscience. 2021;24(12):1711–1720.

18. Wei W, Merkt B, Rotter S. A theory of orientation selectivity emerging from randomly sampling the visual field. bioRxiv. 2022; p. 2022–07.

19. Pinault D. A neurophysiological perspective on a preventive treatment against schizophrenia using transcranial electric stimulation of the corticothalamic pathway. Brain sciences. 2017;7(4):34.

20. Pinault D, Smith Y, Deschênes M. Dendrodendritic and axoaxonic synapses in the thalamic reticular nucleus of the adult rat. Journal of Neuroscience. 1997;17(9):3215–3233.

21. Deschênes M, Madariaga-Domich A, Steriade M. Dendrodendritic synapses in the cat reticularis thalami nucleus: a structural basis for thalamic spindle synchronization. Brain research. 1985;334(1):165–168.

22. Brunel N. Dynamics of sparsely connected networks of excitatory and inhibitory spiking neurons. Journal of computational neuroscience. 2000;8(3):183–208.

23. Braitenberg V, Schüz A. Cortical architectonics. In: Cortex: Statistics and geometry of neuronal connectivity. Springer; 1998. p. 135–137.

24. Gil Z, Connors BW, Amitai Y. Efficacy of thalamocortical and intracortical synaptic connections: quanta, innervation, and reliability. Neuron. 1999;23(2):385–397.

25. Richardson RJ, Blundon JA, Bayazitov IT, Zakharenko SS. Connectivity patterns revealed by mapping of active inputs on dendrites of thalamorecipient neurons in the auditory cortex. Journal of Neuroscience. 2009;29(20):6406–6417.

26. Batschelet E, Batschelet E, Batschelet E, Batschelet E. Circular statistics in biology. vol. 111. Academic press London; 1981.

27. Piscopo DM, El-Danaf RN, Huberman AD, Niell CM. Diverse visual features encoded in mouse lateral geniculate nucleus. Journal of Neuroscience. 2013;33(11):4642–4656.

28. Sadeh S, Cardanobile S, Rotter S. Mean-field analysis of orientation selectivity in inhibition-dominated networks of spiking neurons. SpringerPlus. 2014;3(1):148.

29. Ringach D, Shapley R. Reverse correlation in neurophysiology. Cognitive Science. 2004;28(2):147–166.

30. Deepu R, Spreizer S, Trensch G, Terhorst D, Vennemo SB, Mitchell J, et al. NEST 3.1; 2021. Available from: 10.5281/zenodo.5508805.

31. Spreizer S, Mitchell J, Jordan J, Wybo W, Kurth A, Vennemo SB, et al. NEST 3.3; 2022. Available from: 10.5281/zenodo.6368024.

32. Scholl B, Tan AY, Corey J, Priebe NJ. Emergence of orientation selectivity in the mammalian visual pathway. Journal of Neuroscience. 2013;33(26):10616–10624.

33. Pattadkal JJ, Mato G, van Vreeswijk C, Priebe NJ, Hansel D. Emergent orientation selectivity from random networks in mouse visual cortex. Cell reports. 2018;24(8):2042–2050.

34. Priebe NJ. Mechanisms of orientation selectivity in the primary visual cortex. Annual review of vision science. 2016;2:85–107.

35. Wang W, Andolina IM, Lu Y, Jones HE, Sillito AM. Focal gain control of thalamic visual receptive fields by layer 6 corticothalamic feedback. Cerebral cortex. 2018;28(1):267–280.

36. Pinault D. The thalamic reticular nucleus: structure, function and concept. Brain research reviews. 2004;46(1):1–31.

37. Liu XB, Jones EG. Predominance of corticothalamic synaptic inputs to thalamic reticular nucleus neurons in the rat. Journal of Comparative Neurology. 1999;414(1):67–79.

38. Andolina IM, Jones HE, Wang W, Sillito AM. Corticothalamic feedback enhances stimulus response precision in the visual system. Proceedings of the National Academy of Sciences. 2007;104(5):1685–1690.

39. Sillito AM, Jones HE, Gerstein GL, West DC. Feature-linked synchronization of thalamic relay cell firing induced by feedback from the visual cortex. Nature. 1994;369(6480):479–482.

40. Reinhold K, Resulaj A, Scanziani M. Brain state-dependent modulation of thalamic visual processing by cortico-thalamic feedback. Journal of Neuroscience. 2023;43(9):1540–1554.

41. Jurgens CW, Bell KA, McQuiston AR, Guido W. Optogenetic stimulation of the corticothalamic pathway affects relay cells and GABAergic neurons differently in the mouse visual thalamus. PLOS ONE. 2012;7(9):1–14.

42. Wörgötter F, Eyding D, Macklis JD, Funke K. The influence of the corticothalamic projection on responses in thalamus and cortex. Philosophical Transactions of the Royal Society of London Series B: Biological Sciences. 2002;357(1428):1823–1834.

43. Sherman SM. Tonic and burst firing: dual modes of thalamocortical relay. Trends in neurosciences. 2001;24(2):122–126.

44. Alitto H, Rathbun DL, Vandeleest JJ, Alexander PC, Usrey WM. The augmentation of retinogeniculate communication during thalamic burst mode. Journal of Neuroscience. 2019;39(29):5697–5710.

45. Guido W, Lu SM, Vaughan J, Godwin DW, Sherman SM. Receiver operating characteristic (ROC) analysis of neurons in the cat’s lateral geniculate nucleus during tonic and burst response mode. Visual neuroscience. 1995;12(4):723–741.

46. Béhuret S, Deleuze C, Bal T. Corticothalamic synaptic noise as a mechanism for selective attention in thalamic neurons. Frontiers in Neural Circuits. 2015;9:80.

47. Jahnsen H, Llinas R. Voltage-dependent burst-to-tonic switching of thalamic cell activity: an in vitro study. Archives italiennes de biologie. 1984;122(1):73–82.

48. Denman DJ, Contreras D. Complex effects on in vivo visual responses by specific projections from mouse cortical layer 6 to dorsal lateral geniculate nucleus. Journal of Neuroscience. 2015;35(25):9265–9280.

49. Godwin DW, Vaughan JW, Sherman SM. Metabotropic glutamate receptors switch visual response mode of lateral geniculate nucleus cells from burst to tonic. Journal of neurophysiology. 1996;76(3):1800–1816.

50. Tsumoto T, Creutzfeldt O, Legendy C. Functional organization of the corticofugal system from visual cortex to lateral geniculate nucleus in the cat: With an appendix on geniculo-cortical mono-synaptic connections. Experimental Brain Research. 1978;32:345–364.

51. Cudeiro J, Sillito AM. Spatial frequency tuning of orientation-discontinuity-sensitive corticofugal feedback to the cat lateral geniculate nucleus. The Journal of physiology. 1996;490(2):481–492.

52. Andolina IM, Jones HE, Sillito AM. Effects of cortical feedback on the spatial properties of relay cells in the lateral geniculate nucleus. Journal of neurophysiology. 2013;109(3):889–899.

53. Niell CM, Stryker MP. Highly selective receptive fields in mouse visual cortex. Journal of Neuroscience. 2008;28(30):7520–7536.

54. Binzegger T, Douglas RJ, Martin KA. A quantitative map of the circuit of cat primary visual cortex. Journal of Neuroscience. 2004;24(39):8441–8453.

55. Potjans TC, Diesmann M. The cell-type specific cortical microcircuit: relating structure and activity in a full-scale spiking network model. Cerebral cortex. 2014;24(3):785–806.

56. Merkt B, Schüßler F, Rotter S. Propagation of orientation selectivity in a spiking network model of layered primary visual cortex. PLoS Computational Biology. 2019;15(7):e1007080.

